# SKSR1 identified as key virulence factor in *Cryptosporidium* by genetic crossing

**DOI:** 10.1101/2024.01.29.577707

**Authors:** Wei He, Lianbei Sun, Tianyi Hou, Zuwei Yang, Fuxian Yang, Shengchen Zhang, Tianpeng Wang, Na Li, Yaqiong Guo, L. David Sibley, Yaoyu Feng, Lihua Xiao

**Author notes:** Contributed equally to this work (Wei He, Lianbei Sun, Tianyi Hou). Correspondence (L.X.); (Y.F.); (L.D.S.).

## Abstract

*Cryptosporidium parvum* is a major cause of severe diarrhea. Although isolates of this zoonotic parasite exhibit significant differences in infectivity and virulence, the genetic determinants for these traits are not clear. In this study, we used classical genetics to cross two *C. parvum* isolates of different virulence and used bulked segregant analysis of whole-genome sequence data from the progeny to identify quantitative trait loci (QTL) associated with *Cryptosporidium* infectivity and virulence. Of the 26 genes in three QTL, two had loss-of-function mutations in the low-virulence isolates. Deletion of the *SKSR1* gene or expression of the frame-shift mutant sequence reduced the pathogenicity of infection *in vivo*. SKSR1 is a polymorphic secretory protein expressed in small granules and secreted into the parasite-host interface. These results demonstrate that SKSR1 is an important virulence factor in *Cryptosporidium,* and suggest that this extended family may contribute to pathogenesis.

## Introduction

Cryptosporidiosis is a leading cause of severe diarrhea, leading to death and malnutrition in many children in low- and middle-income countries.^1^ It is also a major cause of food- and waterborne disease in high-income countries.^2^ The disease is particularly severe in young children and immunocompromised individuals, leading to malnutrition and significant mortality.^3^ The pathogenesis of cryptosporidiosis is poorly understood.

Of the more than 20 *Cryptosporidium* species identified in humans, *C. hominis* and *C. parvum* are the dominant species responsible for more than 90% of cryptosporidiosis cases. The two differ significantly in their host range, with the former being predominantly an anthroponotic species and the latter being found in many animals in addition to humans.^4^ Due to the lack of convenient culture and animal models for *C. hominis*, most biological studies of *Cryptosporidium* spp. have been conducted with *C. parvum*.^5^

*C. parvum* isolates vary widely in infectivity and virulence. In experimental infections of healthy adults, three *C. parvum* isolates differed in ID_50_, infection intensity, attack rates, and duration of diarrhea.^6^ Differences in virulence among *C. parvum* isolates have also been observed in experimental infections of mice and young livestock.^7–9^ For example, at the subtype level, IIaA15G2R1 has become the dominant *C. parvum* in both humans and animals in most high-income countries.^4,10^ However, the genetic determinants of *Cryptosporidium* virulence and infectivity are not clear, although the results of comparative genomic analysis have provided some clues.^9^

In *Plasmodium falciparum* and *Toxoplasma gondii*, linkage mapping of genetic crosses of isolates has been effective in the study of virulence factors.^11,12^ It has led to the identification of *P. falciparum* erythrocyte membrane protein 1 (PfEMP1) and several rhoptry proteins as major determinants of virulence in *P. falciparum* and *T. gondii*, respectively.^13,14^ However, traditional linkage analysis requires the comparative analysis of numerous recombinant progeny. In recent years, a more efficient linkage mapping technique, the bulked segregant analysis (BSA), has been used to characterize genetic crosses of *P. falciparum* isolates.^15,16^ It compares the frequencies of phenotypically segregating alleles in pools of recombinant progeny, allowing the identification of SNPs associated with phenotypic traits.^17^

In this report, we describe the identification of genes underlying the virulence differences using BSA of progeny from genetic crosses of two *C. parvum* isolates. The role of candidate genes in infection and virulence was validated using gene depletion and replacement techniques. The results show that the small granule protein SKSR1 is a key virulence factor in *C. parvum*.

## Results

### Coinfection of *C. parvum* isolates of different virulence produces recombinant progeny

We used two *C. parvum* isolates of different GP60 subtypes in our crossing studies, including IIdA20G1-HLJ and IIdA19G1-GD. They differ in infectivity (ID_50_ = 0.6 and 5.1 oocysts, respectively) and infection pattern (IIdA20G1-HLJ produces over 10^7^ OPGs for more than 8 weeks compared to a peak OPG of 10^6^ for less than 2 weeks for IIdA19G1-GD) in interferon-γ knockout (GKO) mice. IIdA20G1-HLJ is highly virulent in this animal model, inducing diarrhea, arched backs, lethargy, reduced weight gain, and over 50% mortality (100% in GKO mice pretreated with antibiotics or treated with paromomycin after infection with transgenic lines carrying the neomycin resistance gene). In contrast, IIdA19G1-GD is avirulent in GKO mice. There are 1263 SNPs between the genomes of the two *C. parvum* isolates.^9^

To create a genetic cross between the virulent IIdA20G1-HLJ and the avirulent IIdA19G1-GD, we endogenously tagged the two isolates with different fluorescent proteins using CRISPR/Cas9. In the IIdA20G1-HLJ isolate, we replaced the uracil phosphoribosyltransferase (*UPRT*) gene in chromosome 1 with a sequence encoding the tdTomato protein. Similarly, in the IIdA19G1-GD isolate, we replaced the thymidine kinase (*TK*) gene in chromosome 5 with a sequence encoding the mNeonGreen protein (Figure S1A). PCR analysis showed correct integration of the replacement cassette (Figure S1B). Fluorescence microscopy of purified oocysts showed that nearly 100% of oocysts from isolates IIdA20G1-HLJ and IIdA19G1-GD were red and green, respectively (Figure S1C).

We performed two genetic crosses of fluorescently tagged IIdA20G1-HLJ and IIdA19G1-GD in GKO mice (Figure 1A). Co-infection of the two isolates resulted in the generation of recombinant oocysts that expressed both red and green color under fluorescence microscopy. We purified F1 progeny oocysts from the intestinal contents of infected mice at DPI 4 (Figure 1B). Approximately 28% of the purified oocysts showed both red and green fluorescence (yellow in merged images), while the remaining oocysts were predominantly red or colorless (Figure 1C). We enriched and collected yellow oocysts by flow cytometric sorting.

**Figure 1.**
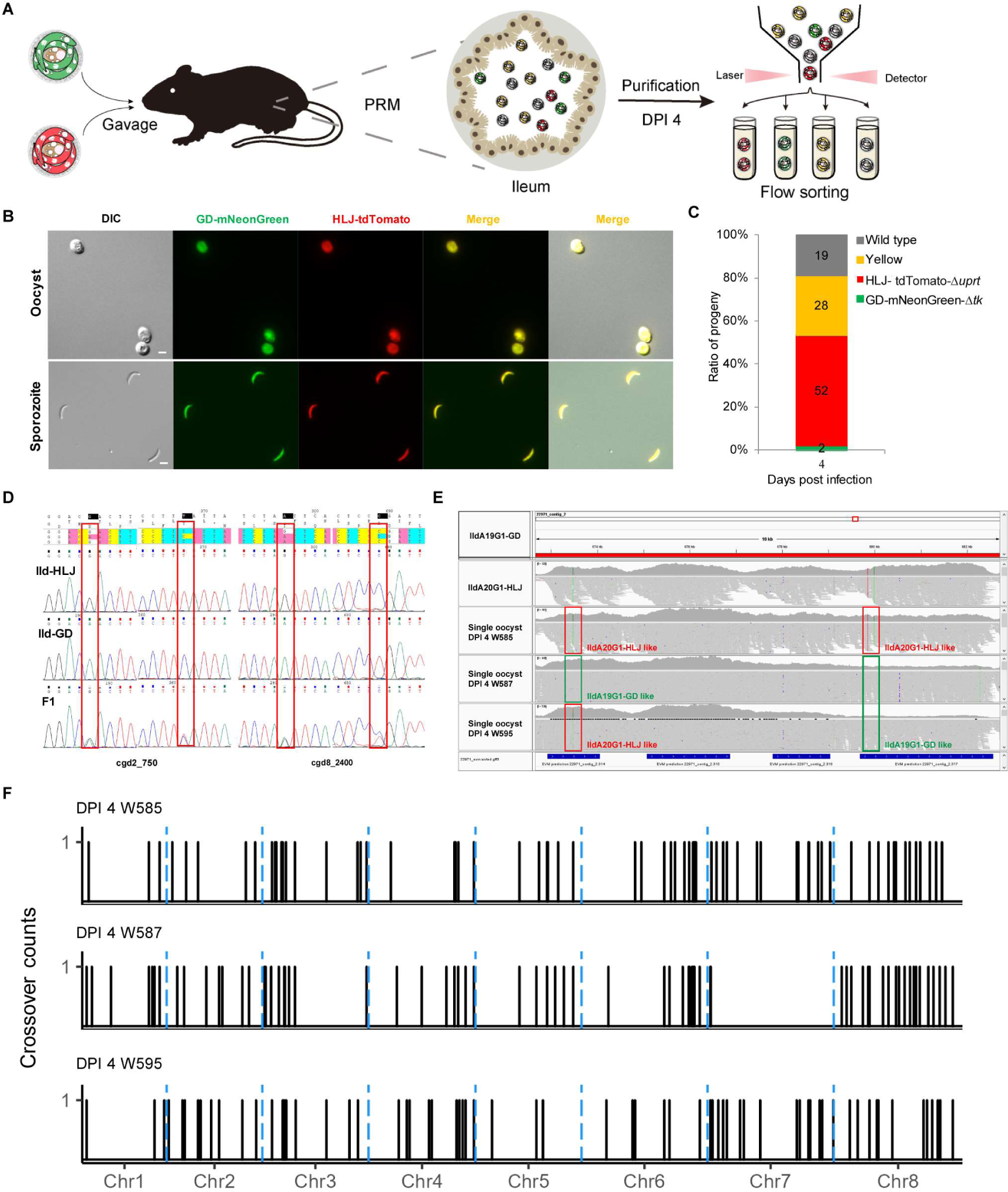
Genetic crosses between IIdA20G1-HLJ and IIdA19G1-GD *in vivo*. (A) Diagram of genetic crosses of virulent IIdA20G1-HLJ and avirulent IIdA19G1-GD in mice and cytometric sorting of progeny. (B) Image of oocysts harvested from intestinal mucosa 4 days after co-infection of oocysts tagged with different fluorescent dyes and the excysted sporozoites. (C) Ratio of oocysts of each color in mouse feces 4 days after coinfection (n = 3). (D) Characterization of cross progeny by sequence analysis of two polymorphic loci. The PCR products of the progeny showed double peaks, indicating that the F1 progeny contained the genomes of two different parents. (E) Confirmation of genetic recombination in individual F1 oocysts by read mapping of whole genome sequences from individual oocysts. Two polymorphic sites relative to the parental sequences on the genomes are marked, and crossover of sequence types is present in oocyst W595. (F) Distribution of recombination events (sequence crossovers) in three F1 oocysts collected 4 days after co-infection. Most sequence crossovers occurred in the subtelomeric regions of the 8 chromosomes.

Based on PCR analysis of polymorphic loci, we confirmed the presence of mixed sequence types in oocysts of dual color (Figure 1D), indicating that the F1 progeny contained genomic sequences from both IIdA20G1-HLJ and IIdA19G1-GD. The recombinant nature of the F1 progeny was confirmed by whole-genome sequencing of individual oocysts, which revealed the presence of mosaic sequences across polymorphic loci in comparative genomic analysis of the data (Figure 1E). Frequent crossovers of the IIdA20G1-HLJ and IIdA19G1-GD sequences were observed along the eight chromosomes of the *C. parvum* genome, suggesting that the yellow oocysts contained pools of recombinant progeny of the genetic cross of IIdA20G1-HLJ and IIdA19G1-GD (Figures 1F and S2).

### Progeny BSA identifies virulence-associated quantitative trait loci

To identify genes associated with virulence, we performed BSA of the progeny from genetic crosses of these two isolates. Fecal samples were collected from GKO mice every 6 days after infection with the F1 progeny for oocyst purification and WGS analysis (Figure 2A). Microscopic examination of oocysts collected at different time points revealed that although only yellow oocysts were used to inoculate mice and paromomycin was used to maintain selection pressure, there was a rapid decrease in the proportion of yellow oocysts during the early course of infection. This change was accompanied by the appearance of red and green oocysts. As the infection progressed, the proportion of green oocysts increased. By DPI 36, there were as many green oocysts as yellow oocysts (Figures 2B and 2C).

**Figure 2.**
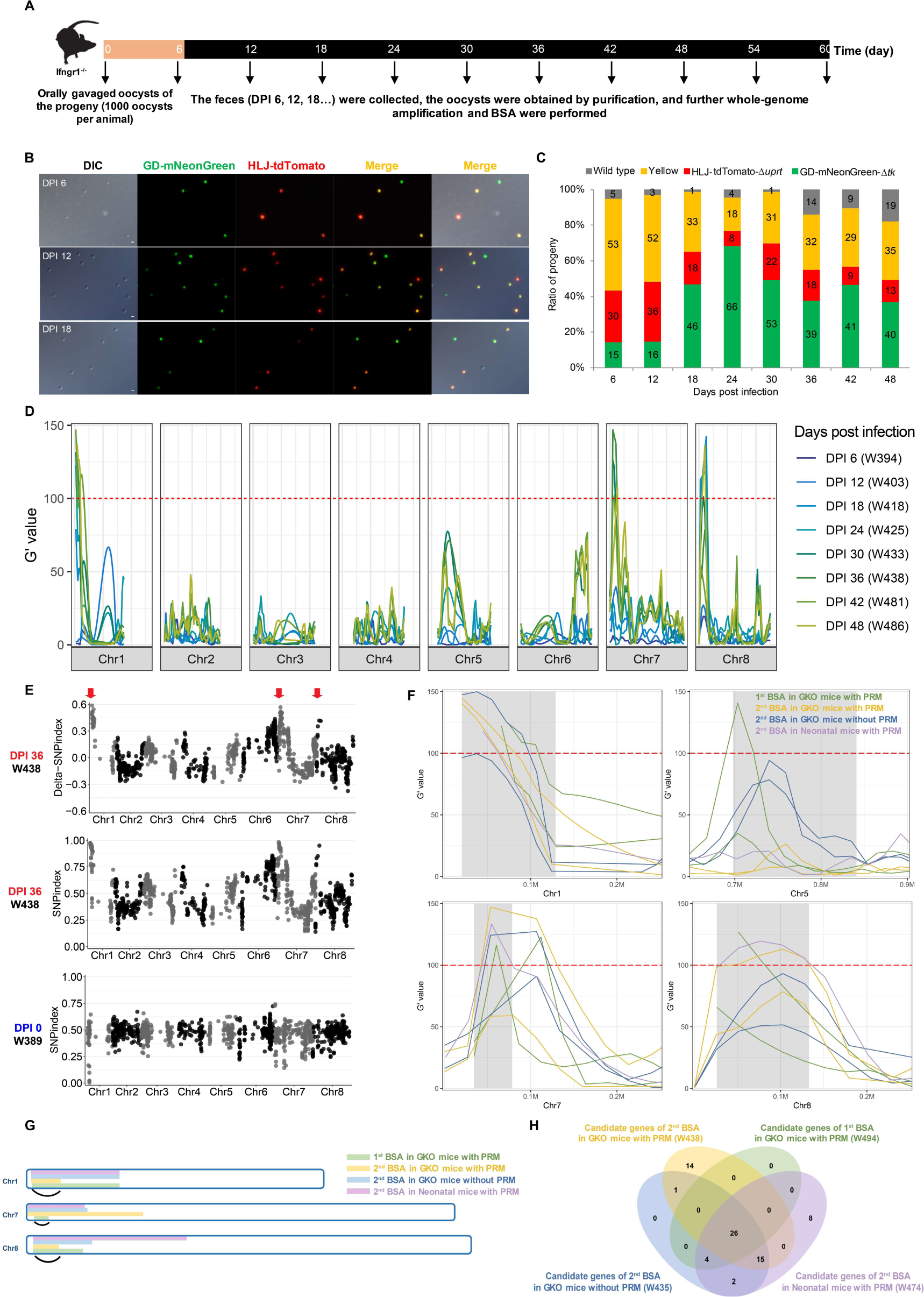
Screening of genes potentially associated with *Cryptosporidium* virulence using the bulked segregant analysis (BSA) (A) Diagram showing the design of the BSA study in GKO mice and the time of sample collection for WGS analysis. (B) Image of purified oocysts at different time points of infection. The first to third panels represent images of purified oocysts at DPI 6, DPI 12, and DPI 18, respectively. Oocyst fluorescence changed significantly after infection, with some oocysts losing fluorescent tags over time. (C) Ratio of parasites with each phenotype at different time points of infection. The fluorescence color in the population changed dynamically during infection. (D) Distribution of G′ values of *Cryptosporidium* genomes collected at different time points of the BSA study. The subtelomeric regions of chromosomes 1, 7, and 8 have high G-statistic values, and there is significant enrichment of sequence alleles at these three for the quantitative trait loci (QTL). (E) Distribution of the SNP index of *Cryptosporidium* genomes collected at different times. The SNP indices in the subtelomeric regions of chromosomes 1, 7, and 8 are close to 1, indicating the enrichment of IIdA20G1-HLJ alleles. (F) Identification of three regions on chromosomes 1, 7, and 8 as the locations of genes underlying the virulence differences between IIdA20G1-HLJ and IIdA19G1-GD, based on a 95% confidence interval. (G and H) Identification of 26 candidate genes associated with virulence using physical and Venn plots.

G-statistical analysis of the WGS data was used to identify SNPs enriched in recombinant *C. parvum* progeny over the course of infection. Gradual increases in G-statistic values were observed in the subtelomeric regions of chromosomes 1, 7, and 8 as infection with the F1 progeny progressed, suggesting an increased frequency of some alleles in these regions (Figure 2D). In addition, the SNP index in these regions was close to 1, suggesting an enrichment of IIdA20G1-HLJ alleles (Figure 2E). In total, we performed four infection studies with two genetic crosses of these two isolates in GKO and neonatal mice with and without the use of paromomycin drug pressure (Figures S3-S5). BSA of WGS data collected at different time points in the infection courses identified similar enrichment of IIdA20G1-HLJ alleles in three regions on chromosomes 1, 7, and 8 (Figure 2F). However, the use of no paromomycin selection during F1 infection of GKO mice identified an additional site of enrichment of IIdA20G1-HLJ alleles on chromosome 5 (Figure S4). In total, the three quantitative trait loci (QTL) were shared by the results of all four BSAs and contained 26 genes on chromosomes 1, 7, and 8. They were associated with the virulence difference between IIdA20G1-HLJ and IIdA19G1-GD (Figures 2G and 2H).

### Candidate QTL genes associated with *C. parvum* virulence

The 26 genes in the three QTL regions were closely examined for sequence differences between IIdA20G1-HLJ and IIdA19G1-GD and the impact levels of these sequence variations (Table S1). The analysis led to the identification of cgd1_140 (*SKSR1*), CPCDC_7g4512, and cgd8_550 as potential determinants of virulence differences between IIdA20G1-HLJ and IIdA19G1-GD. In addition to being polymorphic, these three genes had nucleotide insertions or substitutions that resulted in the formation of stop codons, which are likely to significantly affect the function of the proteins they encode (Figure 3A). Among the three above genes and a paralog (CPCDC_7g4512) of the CPCDC_7g4511 gene (cgd5_4510 in IOWA-II), there was a strong linkage disequilibrium (LD) in sequence between *SKSR1*, CPCDC_7g4511, and CPCDC_7g4512. In contrast, cgd8_550 had a low LD with them (Figures 3B and S6).

**Figure 3.**
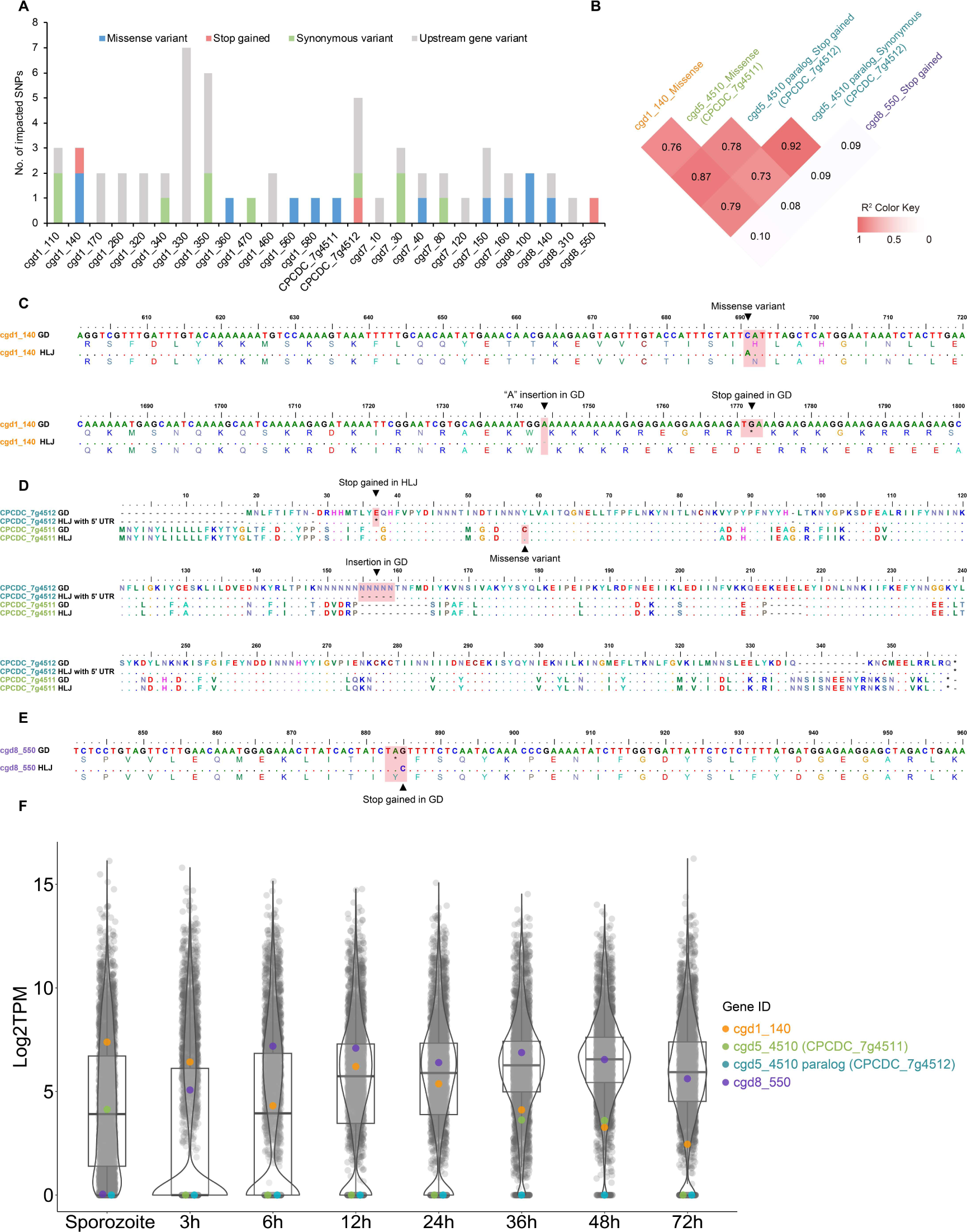
Sequence and expression characteristics of candidate *Cryptosporidium* virulence genes. (A) Distribution of different types of SNPs in 26 genes. (B) Linkage disequilibrium in sequences between potential virulence genes. The *SKSR1*, CPCDC_7g4511 and CPCDC_7g4512 genes have high correlation of sequence polymorphism, while cgd8_550 is poorly correlated with them. (C) Alignment of the partial nucleotide and amino acid sequences of SKSR1-HLJ and SKSR1-GD. Insertion of base A (the first arrowhead) leads to premature termination of SKSR1-GD transcription (the second arrowhead). (D) Amino acid sequence alignment of the partial CPCDC_7g4511 and CPCDC_7g4512 genes. The arrowheads indicate the positions of the stop codon mutation in CPCDC_7g4512 and the missense mutation in CPCDC_7g4511. (E) Alignment of the partial nucleotide sequences and amino acid sequences of the cgd8_550 gene. The arrowhead indicates the position of the base mutation causing a termination codon in cgd8_550-GD. (F) Violin plots of the expression of potential virulence genes. CPCDC_7g4512 is not expressed throughout the life cycle of *Cryptosporidium*, showing the absence of the CPCDC_7g4512 gene in all stages of the life cycle.

Among the three QTL genes, *SKSR1*-GD had an A-base insertion after nucleotide 1753 compared to *SKSR1*-HLJ, resulting in the premature termination of *SKSR1*-GD transcription (Figure 3C). In contrast, the CPCDC_7g4512 gene in the virulent IIdA20G1-HLJ had a G to T substitution, resulting in the formation of a stop codon 51 bp after the start codon. The CPCDC_7g4512 gene had high sequence identity to CPCDC_7g4511, which differed by one nucleotide between IIdA20G1-HLJ and IIdA19G1-GD and had no stop codon-generating nucleotide substitution (Table S1 and Figure 3D). Similarly, there was only one nucleotide difference in the cgd8_550 gene between IIdA20G1-HLJ and IIdA19G1-GD, which resulted in the formation of a termination codon in cgd8_550-GD (Figure 3E).

Since CPCDC_7g4512 is a recently predicted gene, we analyzed its expression level throughout *C. parvum* development. RNA-seq analysis of the transcriptome across all developmental stages of IIdA20G1-HLJ in HCT-8 cultures and sporozoites showed no evidence of CPCDC_7g4512 expression. In contrast, its paralog CPCDC_7g4511 showed modest expression in sporozoites and at 36 and 48 h in culture (Figure 3F). This analysis also showed that although the cgd8_550 gene was not expressed in sporozoites, its expression gradually increased in HCT-8 culture and remained at high levels during 6-48 h. The expression of the *SKSR1* gene was high in sporozoites and at 3, 12 and 24 h in culture (Figure 3F). These data suggested that the *SKSR1* and cgd8_550 genes could be potential determinants of the virulence difference between IIdA20G1-HLJ and IIdA19G1-GD.

### Protein encoded by cgd8_550 is not a virulence factor

We investigated the expression and function of the hypothetical protein encoded by the cgd8_550 gene using genetic manipulation tools. We generated cgd8_550-tagged (cgd8_550-3HA), A-base mutant (cgd8_550^m(G–C)^-3HA), and gene deletion (*Δcgd8_550*) lines of the virulent IIdA20G1-HLJ strain using CRISPR/Cas9 (Figures S7A, S7C and S8A). PCR and fluorescence analysis confirmed the successful deletion of the *cgd8_550* gene in the *Δcgd8_550* line, with oocysts from this line being green due to the replacement of the gene with the mNeonGreen sequence (Figure S7E). IFA analysis of the cgd8_550-3HA line revealed its high expression in trophozoites and meronts. Its expression was much lower in female gametes and male gamonts and absent in free sporozoites and merozoites (Figure S9A). We investigated the role of cgd8_550 in *Cryptosporidium* growth and host pathogenicity using HCT-8 culture and GKO mouse models. Deletion of the cgd8_550 gene did not significantly affect *C. parvum* growth *in vitro* and *in vivo* (Figures S9B and S9C), and GKO mice infected with the cgd8_550-3HA and *Δcgd8_550* lines had similar weight gain and survival (Figures S9D and S9E).

### SKSR1 is a small granule protein encoded by a variant subtelomeric gene

The cgd1_140 gene is a subtelomeric gene on chromosome 1, encoding the *Cryptosporidium*-specific secretory protein SKSR1. Comparison of cgd1_140 sequences from *C. parvum* showed that the gene is polymorphic, with 21-23 SNPs between IIa and IId subtypes in the 2892 bp coding sequence. Within IId subtypes, the A insert between NT1743 and NT1744 was seen in all IIdA19G1 isolates, but was absent from the sequences of other IId isolates. This insert was also not detected in the IIa and IIc isolates analyzed. The sequence polymorphism resulted in the formation of two subclades within the IId cluster and three subclades within the IIa cluster (Figure 4A).

**Figure 4.**
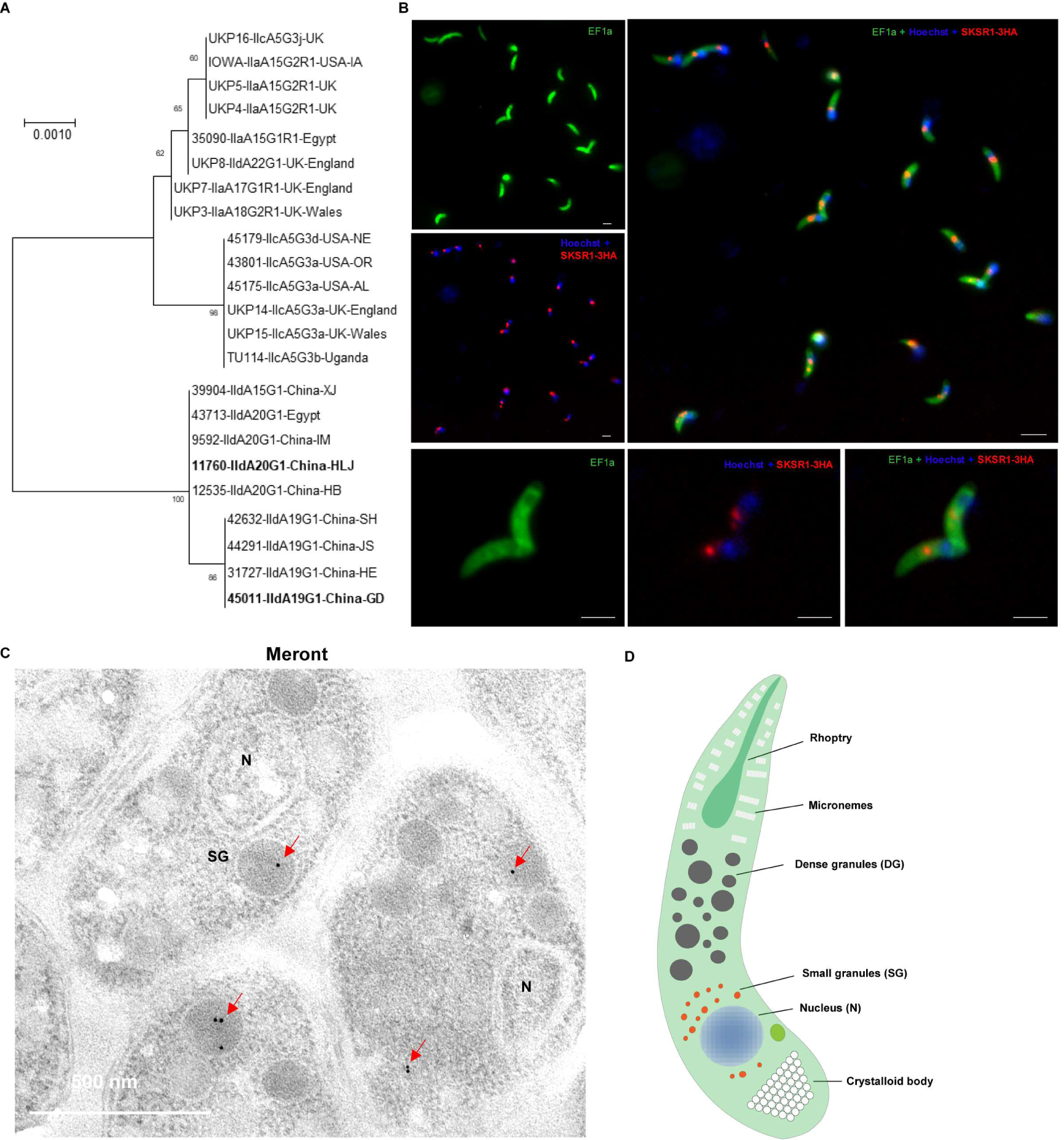
Sequence diversity and expression localization of SKSR1. (A) Sequence diversity and phylogenetic relationship of the *SKSR1* gene in *C. parvum*. The sequence polymorphism leads to the formation of two subclades within the IId cluster and three subclades within the IIa cluster. (B) Localization of SKSR1 expression in sporozoites as revealed by endogenously tagging the gene. SKSR1 is predominantly located near the parasite nucleus. Scale bar: 2 µm. (C) Subcellular localization of SKSR1 by immunoelectron microscopy. SKSR1 is localized to the small granules near parasite nuclei. (D) Schematic representation of the subcellular structure of *Cryptosporidium*.

We generated SKSR1-tagged (SKSR1-3HA), A-base inserted mutant (SKSR1^m(A+)^-3HA), and SKSR1 deletion (*Δsksr1*) lines of the IIdA20G1-HLJ isolate using CRISPR/Cas9 (Figures S7B, S7D and S8B). Fecal luciferase monitoring of mice infected with these lines and PCR analysis of purified oocysts confirmed successful integration of the replacement templates (Figures S7D and S10A). IFA analysis revealed high levels of SKSR1 expression near the nucleus of sporozoites and merozoites (Figures 4B and S10B). SKSR1 expression was significantly lower in female gametes and male gamonts (Figure S10B). We performed immunoelectron microscopy (IEM) of the ileal tissue of mice infected with the SKSR1-3HA line, and the results show the accumulation of gold particles in small vesicles near the nucleus that match the location, shape, and size of small granules (mean = 102 ± 35.2 nm; Figures 4C, 4D, S11A and S11B) rather than canonical dense granules.^33^ In addition, SKSR1 was detected at low levels on the parasite surface and in the feeder organelle (Figure S11C). In the SKSR1^m(A+)^-3HA line, due to the insertion of an A in the *SKSR1* gene, there was no detection of SKSR1 in IFA analysis of all developmental stages (Figure S10B). This was confirmed by Western blot analysis, showing the expression of SKSR1-3HA but not SKSR1^m(A+)^-3HA (Figure S10C). Fluorescence analysis further confirmed the successful deletion of the *SKSR1* gene in the *Δsksr1* line, with oocysts from this line being red due to the replacement of the gene with the tdTomato sequence (Figure S7F).

### SKSR1 is a virulence factor in *C. parvum*

We investigated the role of SKSR1 in *C. parvum* fitness and virulence using *in vitro* and *in vivo* infection models. Compared to the SKSR1-3HA line, deletion of the *SKSR1* gene (*Δsksr1*) significantly reduced *C. parvum* growth *in vitro*. However, the A base insertion in the gene (SKSR1^m(+A)^-3HA) had no apparent effect on parasite growth (Figure 5A).

**Figure 5.**
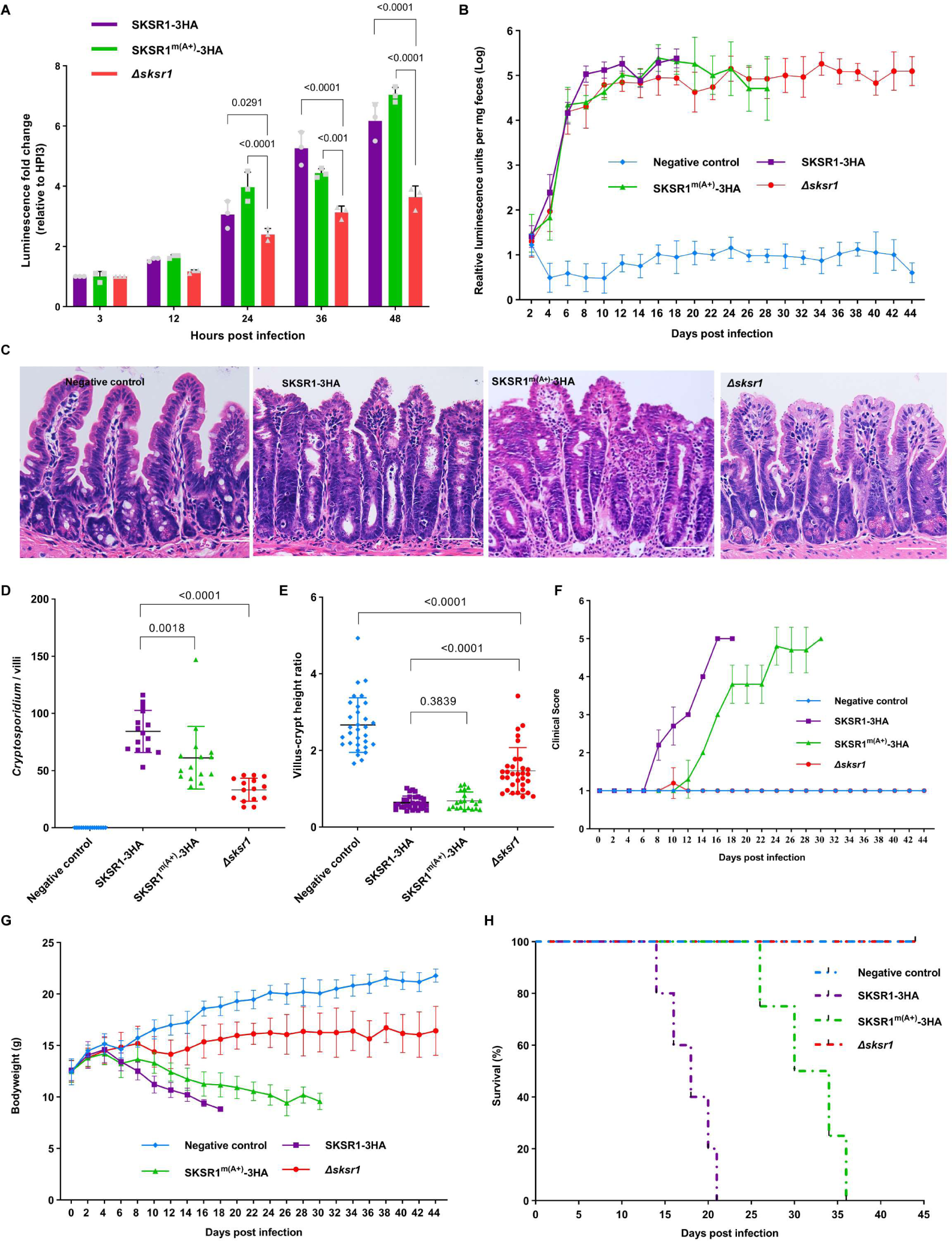
Involvement of SKSR1 in the virulence of *Cryptosporidium parvum*. (A) Growth pattern of *Δsksr1*, SKSR1-3HA, and SKSR1^m(+A)^-3HA lines in *in vitro* culture of HCT-8 cells. The *Δsksr1* line had significantly slower growth at 24 h and 48 h post infection (HPI). (B) Oocyst shedding patterns of *Δsksr1*, SKSR1-3HA, and SKSR1^m(+A)^-3HA lines in GKO mice. Mice (n = 6 per group) were infected with 10,000 oocysts. The oocyst shedding of both *Δsksr1* and SKSR1^m(+A)^-3HA lines was lower than that of the SKSR1-3HA line during the early stages of infection (DPI 8-10). The difference between *Δsksr1* and SKSR1-3HA lines was maintained until the death of all animals in the former group (*p* = 0.0003 at DPI 18). (C) Hematoxylin and eosin (H&E) microscopic images of the ileum of GKO mice infected with transgenic lines compared to the uninfected control. Mice were infected with 10,000 oocysts, killed at DPI 14 and examined for pathological changes during the course of infection. (D) Parasite load on the villi of the small intestine of infected mice. Data are number of parasites per villus under H&E microscopy (*p* < 0.0001 and *p* = 0.0018 for *Δsksr1* and SKSR1^m(+A)^-3HA lines, respectively). (E) Villus length and crypt depth ratio of the ileum from uninfected and infected mice (*p* < 0.0001 and *p* = 0.3839 for *Δsksr1* and SKSR1^m(+A)^-3HA lines, respectively). (F) Differences in clinical score between groups. A higher score indicates a worse condition of the mice during the course of infection. (G) Changes in body weight during the course of infection with different lines. There was a significant difference in body weight between the *Δsksr1* and SKSR1-3HA groups from DPI 8 (*p* = 0.0002), with SKSR1-3HA showing the most significant reduction in body weight. (H) Survival curves of infected mice. The time of mouse death in the SKSR1-3HA group was significantly different from that in the *Δsksr1* and SKSR1^m(+A)^-3HA groups (*p* = 0.0018 and *p* = 0.0049 for the *Δsksr1* and SKSR1^m(+A)^-3HA lines, respectively).

We used the GKO mouse model to examine the differences in growth and host pathogenicity between the *Δsksr1*, SKSR1-3HA, and SKSR1^m(+A)^-3HA lines. During the early stage of infection (DPI 8 - DPI 12), there was no significant difference in oocyst shedding between the *Δsksr1* and SKSR1^m(+A)^-3HA groups. However, the oocyst shedding intensity of these two groups was significantly lower than that of the SKSR1-3HA group. After DPI 12, oocyst shedding in the *Δsksr1* group remained lower than that in the SKSR1-3HA group, although there was no significant difference between the SKSR1-3HA and SKSR1^m(+A)^-3HA groups (Figure 5B). When the intestinal tissues of infected mice were analyzed, the parasite load on the villus surface was significantly different between the SKSR1-3HA and *Δsksr1* groups, with the SKSR1^m(+A)^-3HA group having a parasite load between the two groups (Figures 5C and 5D). SKSR1-3HA and SKSR1^m(+A)^-3HA caused severe intestinal damage, whereas *Δsksr1* caused only mild villous atrophy (Figure 5C). Mice infected with the *Δsksr1* line had a higher villus height to crypt depth ratio than those infected with the SKSR1-3HA and SKSR1^m(+A)^-3HA lines (Figure 5E).

We used a scoring system to rank the severity of clinical signs of infected mice in each group, with higher scores representing worse conditions (Figure 5F). Mice in the SKSR1-3HA group showed significant clinical signs and weight loss at DPI 8, with all animals being dead by DPI 21, while mice in the SKSR1^m(+A)^-3HA group had significantly delayed onset of clinical signs and weight loss (starting at DPI 12), and showed less weight loss than those in the SKSR1-3HA group. The onset of death in the SKSR1^m(+A)^-3HA group occurred at DPI 26, and all mice died by DPI 36. In contrast, *Δsksr1* mice show no clinical signs and weight loss throughout the course of infection, and no mice in these groups died by the end of the infection study at DPI 44 (Figures 5G and 5H).

## Discussion

Virulence factors play a key role in the pathogenesis of microbial pathogens.^18^ They are often involved in the initial pathogen-host interactions, induce cellular damage, or alter host cellular responses. Different pathogens use different strategies to mediate virulence. In apicomplexan parasites, *T. gondii* uses a family of rhoptry-secreted polymorphic kinases as key virulence factors.^12^ They are injected into host cells and mediate immune evasion and other changes in host cell signaling.^19^ In contrast, *P. falciparum* uses a family of variant PfEMP1 surface antigens encoded by the subtelomeric *var* genes. In addition to inducing immune evasion, these proteins mediate the adhesion of infected erythrocytes to endothelial cells in various organs, leading to different types of severe malaria.^20^ Linkage mapping of genetic crosses of isolates of different virulence has played a major role in the identification of virulence factors in these apicomplexan parasites.^11,12^

In the present study, we have successfully used linkage mapping of genetic crosses to identify one virulence factor in *C. parvum*. We have generated genetic crosses of fluorescently tagged *C. parvum* isolates IIdA20G1-HLJ and IIdA19G1-GD, which differ significantly in infectivity, infection pattern, and virulence in GKO mice.^9^ Linkage mapping using BSA of genomes collected over the course of infection with F1 progeny has identified three QTL containing 26 genes that potentially control the virulence difference between the two isolates. Reverse genetic studies of the two best candidates have confirmed the involvement of SKSR1, a secretory small granule protein encoded by a polymorphic subtelomeric gene, in *C. parvum* virulence.

The generation of genetic crosses of *C. parvum* is greatly facilitated by the endogenous fluorescent tagging of isolates. Previous attempts at genetic crosses of *Cryptosporidium* isolates have been hampered by the lack of genetic tagging tools, making it difficult to isolate recombinant progeny difficult without fluorescent tagging and drug selection.^21,22^ Recent advances in genetic manipulation of *Cryptosporidium* spp. allow for endogenous tagging of isolates with fluorescent proteins and incorporation of selectable markers and luciferase sequences for easy detection and enrichment of recombinant progeny.^23^ This approach has been successfully used *in vitro* to generate genetic crosses of a *C. parvum* isolate tagged with two different fluorescent dyes.^24^ Research has also shown that genetic crosses between species are possible. Crosses between *C. parvum* and *C. tyzzeri* have resulted in progeny with a recombinant genome derived from both species that continues to reproduce vigorously sexually.^25^ Despite the fact that these two species hybridize, the extent of recombination on most chromosomes is uncertain, limiting the ability to identify genes related to infectivity or host range. In the present study, we have tagged two *C. parvum* isolates of different virulence and generated genetic crosses in mice, and purified the recombinant progeny using flow cytometric sorting. WGS analysis of individual oocysts has confirmed the presence of frequent crossovers in the genomic sequences of the progeny, thus facilitating genetic mapping and gene identification.

The identification of virulence determinants in *Cryptosporidium* spp. in the present study is achieved by a new approach to perform linkage mapping of recombinant progeny using BSA of WGS data. The technique examines genomic changes during the course of infection using pools of progeny from genetic crosses. Alleles enriched in late infection are identified by read mapping of WGS data.^17^ This eliminates the need to clone the progeny of genetic crosses, which is difficult to perform for *Cryptosporidium* spp. due to the lack of effective culture systems and the high ID_50_ of isolates in laboratory animal models. This strategy has been used in combination with crosses to map genes associated with drug resistance, virulence and other phenotypic traits in *P. falciparum*. ^15,17^ In the present study, by BSA of 68 sets of WGS data collected from GKO and neonatal mice infected with the F1 progeny of two genetic crosses of *C. parvum* isolates of low and high virulence, we have identified the evolution toward the more virulent parent as the infection persisted. This is consistent with the observation in a previous study of genetic crosses of two *C. parvum* isolates with different host preferences.^22^ Using this approach, we have identified three QTL containing 26 genes on chromosomes 1, 7, and 8 that are potentially involved in the virulence difference between the two *C. parvum* isolates studied.

Most of the 26 candidate genes have only a small number of SNPs that are likely to have little effect on the functions of the proteins they encode, and thus are unlikely to contribute significantly to the difference in virulence between the two *C. parvum* isolates studied. These include synonymous substitutions and SNPs in non-coding regions. However, three of these genes, including cgd1_140 (*SKSR1*), CPCDC_7g4512, and cgd8_550, have premature stop codons due to single base insertions/deletions and loss-of-function SNPs. Among them, CPCDC_7g4512 has the deleterious nucleotide substitution in the highly virulent isolate IIdA20G1-HLJ. Since this newly predicted gene is not expressed at different developmental stages, CPCDC_7g4512 is unlikely to be involved in virulence. Its identification among the QTL genes is probably the result of the presence of linkage disequilibrium in the sequences between *SKSR1* and CPCDC_7g4512. This may be due to genetic hitchhiking associated with the fixation of beneficial mutations.^26,27^ In contrast, cgd1_140 (*SKSR1*) and cgd8_550 have premature stop codons in the low virulence isolate IIdA19G1-GD, suggesting that they are candidate virulence determinants.

Results from gene tagging and deletion studies suggest that the cgd8_550 gene is unlikely to be involved in virulence. The sequence characteristics of the gene are not related to those of other QTL identified in the study. In addition, the protein it encodes is a cytosolic protein that is predominantly expressed in the asexual stages rather than in the invasive sporozoites and merozoites, suggesting that it is unlikely to interact with host cells as expected for known virulence factors in apicomplexans. Indeed, depletion of the gene in the virulent IIdA20G1-HLJ has minimal effect on the growth and pathogenicity of the isolate.

Studies using genetically modified lines of IIdA20G1-HLJ have confirmed the involvement of SKSR1 in *C. parvum* virulence. Deletion of the *SKSR1* gene significantly reduced *C. parvum* growth *in vitro*, and attenuated parasite virulence *in vivo*. In particular, mice infected with the *Δsksr1* line had significantly reduced infection intensity and no weight loss and mortality. More importantly, replacement of the *SKSR1* gene in IIdA20G1-HLJ with IIdA19G1-GD also reduced the infection intensity and pathogenicity of the wild isolate, resulting in reduced weight loss and improved survival of infected mice. SKSR1 shares some characteristics of virulence factors in other apicomplexans, including being a secretory protein encoded by a polymorphic subtelomeric gene, being a member of a large protein family, and being secreted into the parasite-host interface during invasion and early development.^28,29^ The SKSR1 family is one of a small number of multigene families in *Cryptosporidium* spp. They are secreted proteins encoded by 8-11 subtelomeric genes. These genes are polymorphic within and between *Cryptosporidium* species, and are mainly found in species closely related to the human-pathogenic *C. parvum* and *C. hominis*.^30^ In *C. parvum*, gain and loss of SKSR genes is associated with the host range of isolates.^31^ However, very few studies have investigated the biological function of SKSR proteins. One study genetically tagged SKSR7 and SKSR8, but their locations and functions remain unknown.^32^

The findings of this study may shed light on the biological function of SKSR1. First, we have identified SKSR1 as a member of the secretory proteins in the newly identified small granules (SG). Recently, two SG proteins were identified in *C. parvum* using spatial proteomics. Although the functions of these proteins remain unclear, SG1 has been shown to be secreted into the parasite-host interface soon after invasion and may thus may be involved in host-pathogen interactions.^33^ Second, SKSR1 is secreted into the PVM and feeder organelle, suggesting that it has the potential to manipulate the host cellular signaling. In *T. gondii*, several dense granule proteins are secreted into the PVM to modulate host cellular responses as part of the parasite’s immune evasion.^34^ Therefore, SKSR1 may act as a virulence factor through regulating host responses, a possibility that awaits future study.

## Methods

### Parasite isolates

The IIdA19G1-GD and IIdA20G1-HLJ isolates of *C. parvum* used in this study were originally obtained from dairy calves in Guangdong (GD) and Heilongjiang (HLJ) provinces, China, respectively ^9^ and propagated in interferon-γ knockout (GKO) C57BL/6J mice. *C. parvum* oocysts were purified from the feces of infected mice by sucrose flotation and cesium chloride gradient centrifugation as described.^9^

### Mice

GKO mice were purchased from The Jackson Laboratory (Bar Harbor, USA) and bred in-house at the Laboratory Animal Center of the South China Agricultural University. This study was approved by the Institutional Animal Care and Use Committee of South China Agricultural University. Animals used in the study were housed and handled in accordance with established ethical principles.

### Construction of CRISPR/Cas9 plasmids

Homologous repair templates and CRISPR/Cas9 plasmids were generated as previously described.^24^ The CRISPR/Cas9 plasmids were constructed by inserting an sgRNA targeting the gene of interest into the *C. parvum* Cas9/U6 linear plasmid amplified from pACT1:Cas9-GFP, U6:sgTK using Gibson assembly cloning. To generate the GD-*Δtk*-mNeonGreen and HLJ*-Δuprt*-tdTomato lines, we used mNeonGreen and tdTomato tags to replace GFP and mCh in TK-GFP-Nluc-P2A-neo-TK and UPRT-mCh-Nluc-P2A-neo-UPRT plasmids, respectively. To generate gene knockout lines (*Δsksr1* or *Δcgd8_550*), homology repair fragments were generated by PCR amplification of TK-mNeonGreen-Nluc-P2A-neo-TK and UPRT-tdTomato-Nluc-P2A-neo-UPRT. These fragments were assembled with corresponding regions of the gene of interest, including the 3’ and 5’ UTR sequences. To tag genes with the 3 × HA epitope, we modified the pINS1-3HA-Nluc-P2A-neo^35^ by replacing its *INS1* C-terminal and 3’ UTR sequences with regions of the gene of interest using a four-fragment Gibson assembly. In addition, to prevent erroneous cleavage of the repair plasmid by the CRISPR/Cas9 plasmid, we modified the gRNA or PAM sequence at the 5’ homology arm of the repair plasmid using a two-fragment Gibson assembly.

The primers used in this study are listed in Table S2.

### Generation of transgenic parasites

Transgenic parasites were generated using CRISPR/Cas9 as previously described.^23,36^ Briefly, oocysts were treated with 25% Clorox (1.3% sodium hypochlorite) on ice for 10 minutes and excysted by treatment with sodium taurodeoxycholate at 37°C for 60 minutes. The excysted sporozoites were transfected with an appropriate CRISPR/Cas9 plasmid containing a parasite-specific guide RNA (gRNA) and a targeting plasmid containing the desired genetic manipulation flanked by DNA homologous to the regions surrounding the gRNA cut site. The transfected sporozoites were administered by oral gavage to GKO mice pretreated with oral sodium bicarbonate to neutralize gastric acid. Transgenic parasites in infected mice were selected by continuous administration of 16 g/L paromomycin in the drinking water at 18 hours post infection (HPI 18).

### Measurement of parasite burden by luciferase assay

To quantify transgenic parasites expressing luciferase, fecal pellets from infected mice were weighed, placed in microfuge tubes containing 1 mL luciferase lysis buffer and glass beads, and vortexed for 3 minutes. The tubes were briefly centrifuged at 10,000 × g to pellet debris, and 100 μL of the supernatant was transferred to a 96-well round-bottom plate (Costar, USA). Next, 100 μL of a 1:50 mixture of nanoluciferase substrate:buffer (Promega, USA) was added to each sample, and luminescence was measured using a BioTek Synergy H1 Hybrid plate reader (BioTek, USA).

### PCR analysis of transgenic parasites to confirm gene integration

DNA was extracted from oocysts of transgenic parasites using the QIAGEN DNeasy Blood and Tissue Kit (QIAGEN, Germany). PCR primers were designed to anneal outside the 5′ and 3′ homology arms directing homologous recombination, the luciferase reporter gene and the neomycin selection marker. In addition, primers were designed for the coding and mutant regions of the gene of interest to verify successful knockout and mutation of genes.

### Genetic crossing of isolates with different virulence

Two GKO mice were treated orally with a suspension containing 10,000 each of HLJ-*Δuprt*-tdTomato oocysts and GD-*Δtk*-mNeonGreen oocysts. Infected mice were treated continuously with 16 g/L paromomycin via drinking water at HPI 18. They were monitored for fecal luciferase at DPI 4. If the mice were positive, they were euthanized and the small intestine of the mice was collected. The small intestine was rinsed several times with chilled PBS, and the suspension was filtered through 70 μm and 40 μm filters (Falcon, Cat#352350 and Cat#352350). The filtrate was centrifuged at 10,000 × g for 10 minutes, and the pellet was resuspended in 500 μL PBS for fluorescence microscopy on a BX53 microscope (Olympus, Japan). The oocyst suspension was also analyzed by flow cytometry on a FACSAria III (BD Biosciences, USA), with recombinant oocysts of dual color harvested by flow sorting. Two genetic crosses of the two isolates were performed during this study.

### Isolation of single oocysts for whole genome sequencing

The F1 progeny were diluted to a final concentration of 1 oocyst/μL, and 1 μL of the oocyst suspension was placed on a low adsorption slide for light microscopy. After confirming the presence of a single oocyst, the droplet was transferred to a PCR tube with 2.5 μL PBS form the droplet wash. The harvested single oocyst was repeatedly freeze-thawed in liquid nitrogen and a 55°C water bath to lyse the oocysts. The released genomic DNA was amplified using the REPLI-g Single Cell Kit (QIAGEN) and analyzed by PCR targeting three genes that were polymorphic between the IIdA19G1-GD and IIdA20G1-HLJ isolates. Positive DNA from individual oocysts was sequenced for whole genomes as described below. The PCR sequences used are listed in Table S2.

### Infection of mice with progeny of genetic crosses

In the first infection study, three GKO mice were orally gavaged with 1000 yellow oocysts from the F1 progeny of the genetic cross and treated with paromomycin as described above. Fecal samples were collected from the infected mice every 6 days for 36 days starting at DPI 6. In the second infection study, two infection groups were established with and without paromomycin treatment, and animals (n = 3) in each group were infected with F1 progeny from the second cross. A total of 49 samples were collected at multiple time points of infection as described in the first infection experiment. In the third infection study, one liter of 5-day-old C57BL/6J mice were infected as in the second infection study and 4 samples were collected at DPI 12, DPI 18, DPI 24, and DPI 30. Oocysts in fecal samples collected during the infection studies were purified by cesium chloride and sucrose gradient centrifugation for whole genome sequencing.

### Library preparation and sequencing

Genomic DNA was extracted from the purified oocysts using the Qiagen DNA Mini Kit and amplified using the REPLI-g Midi Kit (QIAGEN). WGA products were purified using the EasyPure PCR Purification Kit (TransGen Biotech, China), and sequenced using Illumina 150 bp paired-end technology on a Novaseq S4 or Hiseq X sequencer to achieve >100-fold coverage of the *Cryptosporidium* genome. In addition to sequencing 12 individual oocysts from the F1 progeny, 68 oocyst pools from the F1 progeny infection studies were whole-genome sequenced during the study.

### Mapping, genotyping and BSA

Fastq files from WGS were checked for quality using FastQC-v0.11.05, and sequence reads were trimmed for poor quality and adapters using Trimmomatics v0.39.^37^ The cleaned reads were aligned to the *C. parvum* IIdA19G1 genome using BWA MEM2 v2.2.^38^ The resulting alignments were sorted and duplicate reads were removed using SAMtools v1.7 and Sambamba v0.8.2.^39,40^

For BSA, the previously described approach was used.^15^ Briefly, variants were called and filtered using BCFtools v1.12^39^ and annotated using SnpEff v4t.^41^ The modules *runQTLseqAnalysis* and *runGprimeAnalysis* in the R package QTLseqr were used to calculate the delta SNP index and G’ values.^42^ Regions with G’ >100 were identified as extreme QTL. Once a QTL was detected, genes within the QTL regions were extracted based on the intersection of the QTL regions obtained from the BSA replicate experiments using jvenn (https://jvenn.toulouse.inra.fr/app/example.html).

### Assessment of transgenic parasite development and immunofluorescence assay *in vitro*

HCT-8 cells were seeded on 48-well plates and grown to 80% confluence. Bleach-treated transgenic parasite oocysts (10,000 oocysts per well) with identical luminescence levels were then used to infect the HCT-8 monolayer for various time periods: 3, 12, 24, 36, and 48 hours. Cultures infected with transgenic oocysts were then washed twice with PBS at HPI 3 and replenished with fresh medium containing 2% fetal bovine serum. At multiple time points, the culture medium was removed from the wells, 100 µL of lysis buffer was added to the wells, and the plate was incubated at 37°C for 10 minutes. After incubation at 37°C for 10 minutes, the lysates were collected by centrifugation at 15,000 × g for 3 minutes and subjected to luciferase assay as described previously.

Coverslips were placed in 24-well plates and seeded with HCT-8 cells. After establishing the *in vitro* infection pattern as described above, the cells were washed with PBS, fixed by incubation with 4% paraformaldehyde, and permeabilized by treatment with 0.5% Triton X-100 for 15 minutes. The coverslips were then blocked with 1% bovine serum albumin (BSA) and incubated with antibodies diluted in the blocking solution, including a rabbit monoclonal antibody against HA (1:800) as primary antibody and goat anti-rabbit polyclonal Alexa Fluor 594 (1:400) as secondary antibody, along with direct staining of parasite stages using Vicia Villosa Lectin (1:1000). Hoechst was used to stain host cell and parasite nuclei. Slides were examined using either a Zeiss Imager M2 microscope or an Olympus BX53 microscope. For immunofluorescence analysis of sporozoites, sporozoites were resuspended in 10 µL PBS and spread on sterile coverslips pretreated with poly-L-lysine. After fixation and permeabilization, they were treated with mouse anti-Cp23 (1:200) or anti-EF1a (1:200) and rabbit monoclonal anti-HA (1:800) as primary antibodies and goat anti-rabbit polyclonal Alexa Fluor 594 (1:400) and goat anti-mouse polyclonal Alexa Fluor 488 (1:400) as secondary antibodies. Each analysis was performed in duplicate for at least two biological replicates.

### Western blot analysis of transgenic parasite oocysts

Purified transgenic oocysts (5 × 10^6^) were treated with bleach, washed with cold PBS, and resuspended in lysis buffer (ThermoFisher Scientific, USA) containing protease inhibitors (Sigma-Aldrich, USA). The mixture was incubated overnight at 4°C, combined with protein loading buffer, and boiled for 10 minutes. Proteins in the lysate were then fractionated by SDS-PAGE and transferred to a PVDF membrane (Merck Millipore, Cat#IPVH00010). After blocking with 1% nonfat milk overnight at 4°C, the membrane was incubated with anti-HA primary antibody (1:2000) for 1 hour, and washed three times with PBS containing 0.1% Tween 20 (PBST). After incubation with HRP-conjugated anti-rabbit IgG (H + L) (1:10,000), washed three times with PBST, and treated with High-sig ECL Western Blotting Substrate, the membrane was analyzed using a Tanon 5200 (Tanon, China). The membrane was further probed with a mouse polyclonal antibody (1:2000) against CP23 of *C. parvum* and HRP-conjugated goat anti-mouse IgG (H + L) (1:10,000).

### Histological and immunoelectron microscopy analysis of infected tissue

For studies of the biological significance of virulence-related genes, one mouse from each experimental group was selected and euthanized on day 14 post infection (DPI) at the peak of oocyst shedding. The small intestine was dissected and washed with cold PBS. The ileum was harvested for conventional hematoxylin-eosin (H&E) and IEM. For H&E microscopy, 30 villi were selected for measurement of villus length and crypt height and calculation of villus length/crypt height ratio. Parasite burden was also determined in 15 intestinal villi. For IEM, rabbit anti-HA (1:20) and goat anti-rabbit IgG conjugated with 10 nm colloidal gold (1:20) were used as primary and secondary antibodies, respectively, as described.^35^ The processed sections were examined on a Talos L120C (ThermoFisher Scientific, USA).

### Assessment of virulence of transgenic *C. parvum* in mice

To assess the biological significance of each candidate gene, GKO mice in each infection group were orally gavaged with 10,000 oocysts of transgenic *C. parvum*, and received paromomycin via drinking water as described above. After infection, fecal luciferase activity and body weight of each mouse were determined every other day, and diarrhea, mortality, and other clinical signs were recorded daily. A scoring system was used to evaluate the severity of the clinical signs in infected mice: 1: mice were in good health and physically active; 2: mice appeared depressed and moved less frequently; 3: mice were depressed, had arched backs, and moved only when touched by hand; 4: mice had hunched position, rough hair, fecal caking, and were immobile even when touched. The difference between groups was evaluated by two-way ANOVA for multiple comparisons. Survival curves were plotted at the end of the infection study, and the difference between groups was evaluated by the log-rank Mantel-Cox test.

### Statistical analysis

GraphPad Prism (https://www.graphpad.com/) was used for all statistical analyses. Unless otherwise noted, Student’s t-test was used to assess differences between two groups, and one-way ANOVA with Tukey’s multiple comparison test was used to assess differences between three or more groups.

### Supplemental material

Supplemental material is available online only.

**Figures S1-S11**, TIF file. **Table S1-S2**, XLSX file

## Acknowledgments

This work was supported in part by the National Natural Science Foundation of China (32030109, 31972697, 31820103014, and 32150710530), Guangdong Major Project of Basic and Applied Basic Research (2020B0301030007), 111 Project (D20008), and Double First-class Discipline Promotion Project (2023B10564003). We thank Jilei Huang, Chuanhe Liu, and Xiaoxian Wu of the Instrumental Analysis & Research Center, South China Agricultural University for assistance with electron microscopy.

## Author contributions

Conceptualization: L.X., Y.F. and L.D.S.; Methodology and investigation: W.H., L.S., T.H., Z.Y., F.Y., S.Z., T.W., N.L., and Y.G.; Formal Analysis: W.H., H.S., and T.H.; Supervision: L.X., Y.F. and L.D.S.; Writing—original draft: W.H., L.X., and Y.F. Writing—review and editing: All authors.

## Data and code availability

All sequence data are deposited in the NCBI Short Read Archive (https://www.ncbi.nlm.nih.gov/sra/) under the BioProject accession number PRJNA1069297. All data needed to evaluate the conclusions in the paper are present in the paper and/or the Supplementary Materials. This paper does not report original code.

---

**Figure S1.**
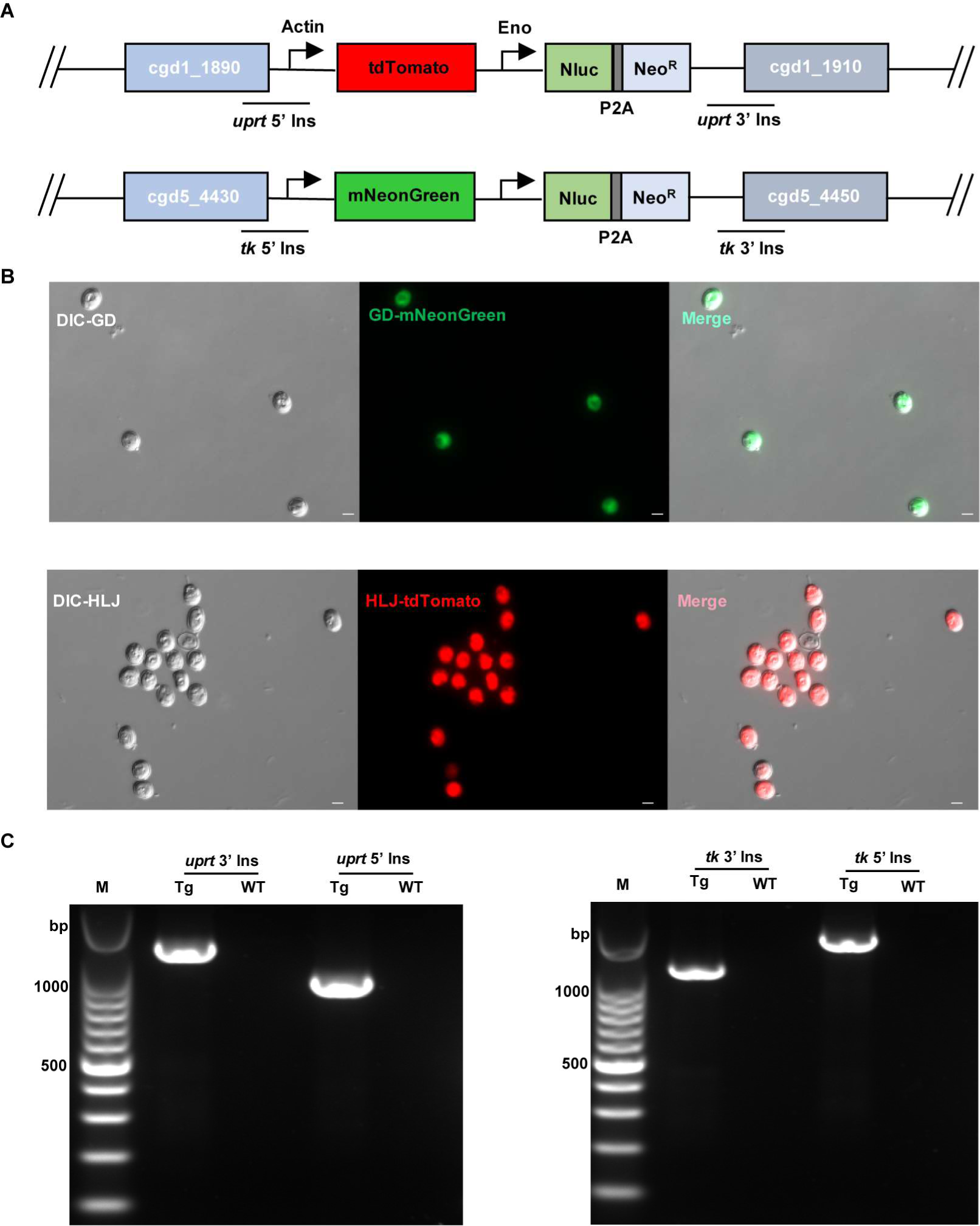
Genetic tagging of two *Cryptosporidium parvum* isolates with different virulence. (A) Illustration of the targeting constructs designed to replace the endogenous *UPRT* (cgd1_1900) and *TK* (cgd5_4440) loci with the tdTomato/mNeonGreen and Nluc-P2A-NeoR cassette, respectively. (B) Image of oocysts purified from fecal samples of GKO mice infected with IIdA19G1-GD transfected with mNeonGreen (GD-mNeonGreen) and IIdA20G1-HLJ transfected with tdTomato (HLJ-tdTomato). (C) PCR confirmation of the correct integration of the tagging constructs in GD-mNeonGreen and HLJ-tdTomato oocysts purified from fecal samples of infected mice.

**Figure S2.**
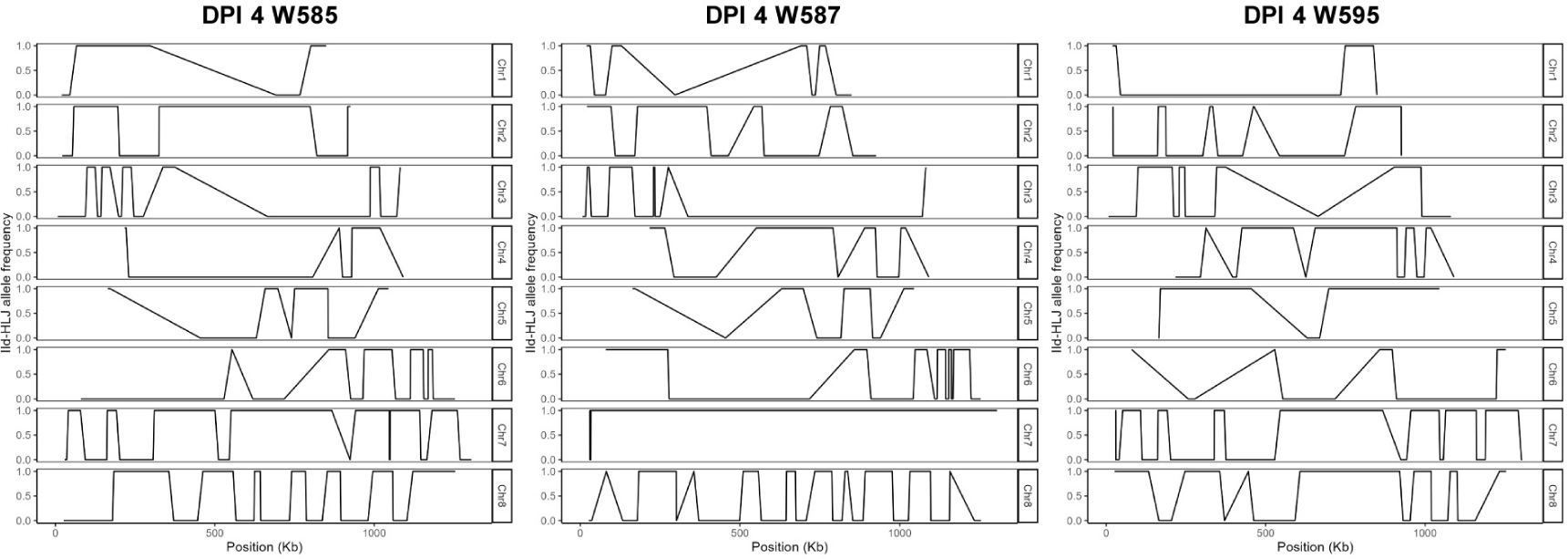
Allele frequency along the 8 chromosomes according of whole-genome sequencing of 3 oocysts of the F1 progeny of genetic crossing of IIdA19G1-GD and IIdA20G1-HLJ.

**Figure S3.**
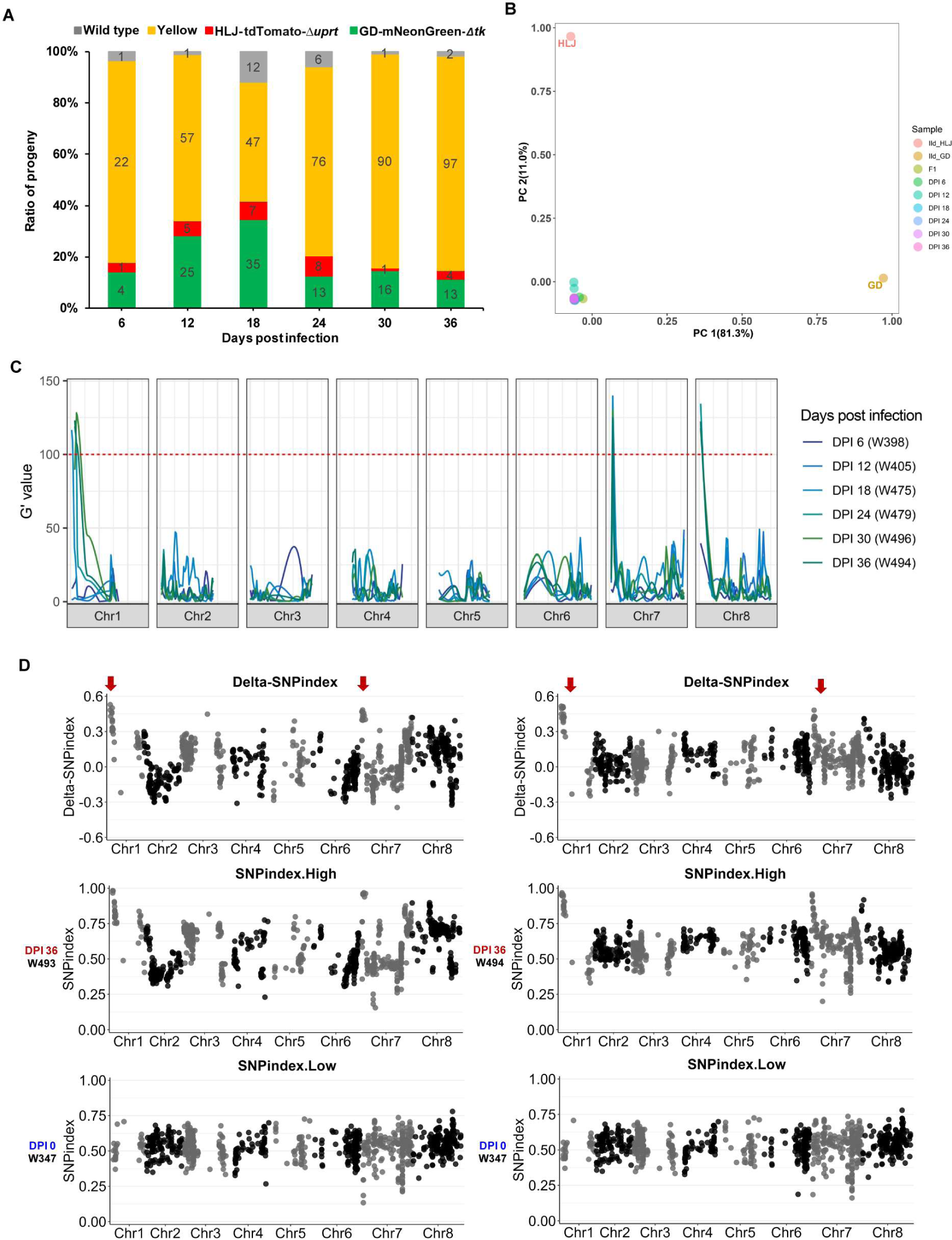
Initial screening of genes potentially associated with *Cryptosporidium* virulence using bulked segregant analysis (BSA) of parasites collected from GKO mice infected with F1 progeny from the first cross of IIdA19G1-GD and IIdA20G1-HLJ. Paromomycin was used throughout the infection. (A) Ratio of oocysts of different colors at different time points of infection. (B) Principal component analysis (PCA) of WGS data from oocysts collected at different times after infection, where PC1 and PC2 account for variability among samples. (C) Distribution of G′ values of *Cryptosporidium* genomes collected at different times of the BSA study. The subtelomeric regions of chromosomes 1, 7, and 8 have high G-statistic values, indicating a significant enrichment of sequence alleles in these regions. (D) Distribution of the SNP index of two samples collected at DPI 36. The SNP indices in the subtelomeric regions of chromosomes 1 and 7 are close to 1, indicating the enrichment of IIdA20G1-HLJ alleles.

**Figure S4.**
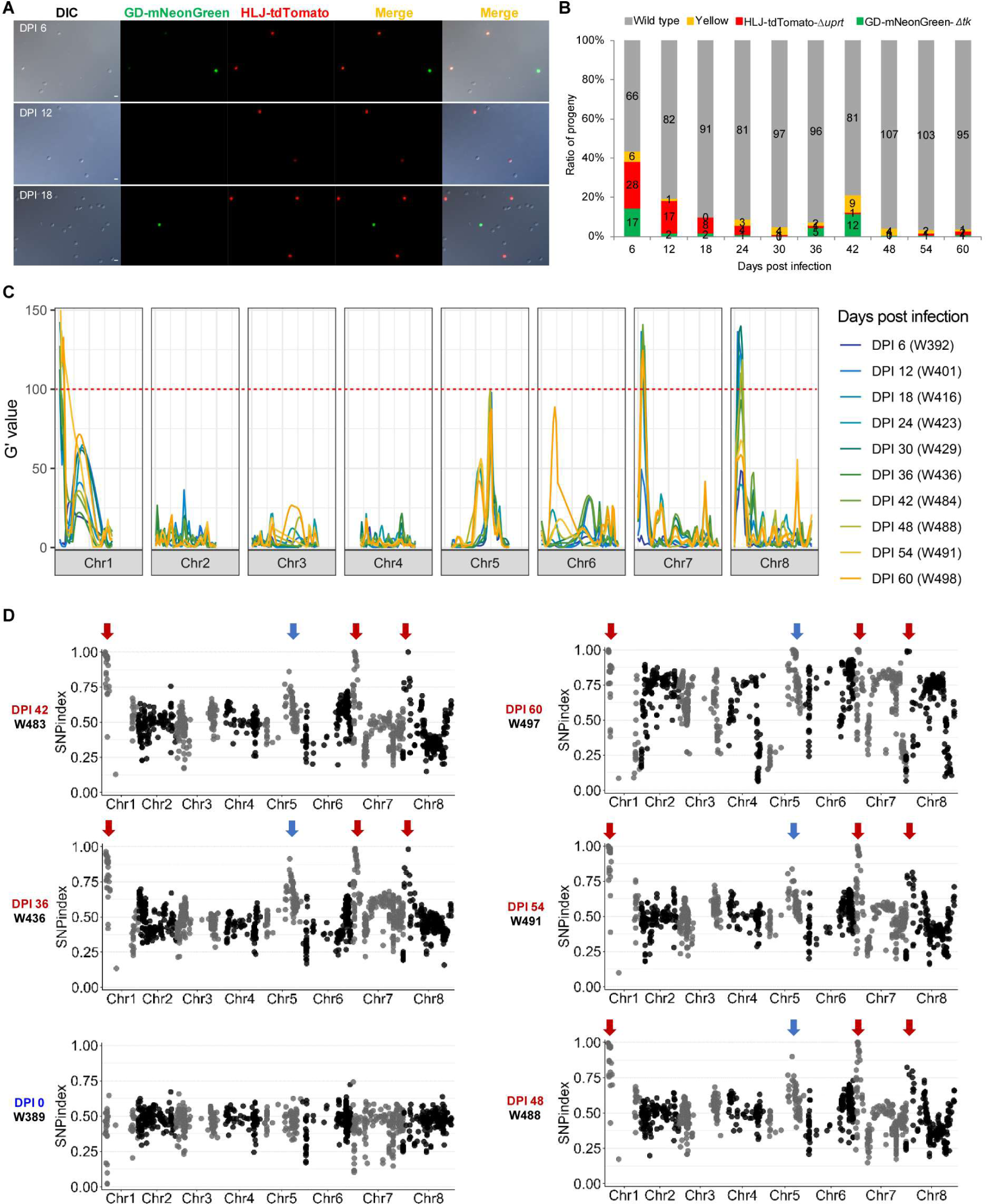
Second screening of genes potentially associated with *Cryptosporidium* virulence using BSA in GKO mice infected with F1 progeny of the second cross of IIdA19G1-GD and IIdA20G1-HLJ treated without paromomycin throughout the infection. (A) Image of purified oocysts at different time points of infection. Fluorescence of oocysts was progressively lost during infection, The first to the third panels represent images of purified oocysts at DPI 6, DPI 12, and DPI18, respectively. (B) The ratio of oocysts of different colors at different time points of infection (n = 3). (C) Distribution of G-statistic values of *Cryptosporidium* genomes collected at different time points of the BSA study. The subtelomeric regions of chromosomes 1, 7, and 8 have high G-statistic values, indicating a significant enrichment of sequence alleles in these regions. (D) Distribution of the SNP index of the samples collected throughout the infection. The SNP indices subtelomeric regions of chromosomes 1, 5, 7, and 8 are close to 1, indicating enrichment of IIdA20G1-HLJ alleles. Some regions of chromosome 5 were enriched compared to the paromomycin-treated group, possibly related to drug screening.

**Figure S5.**
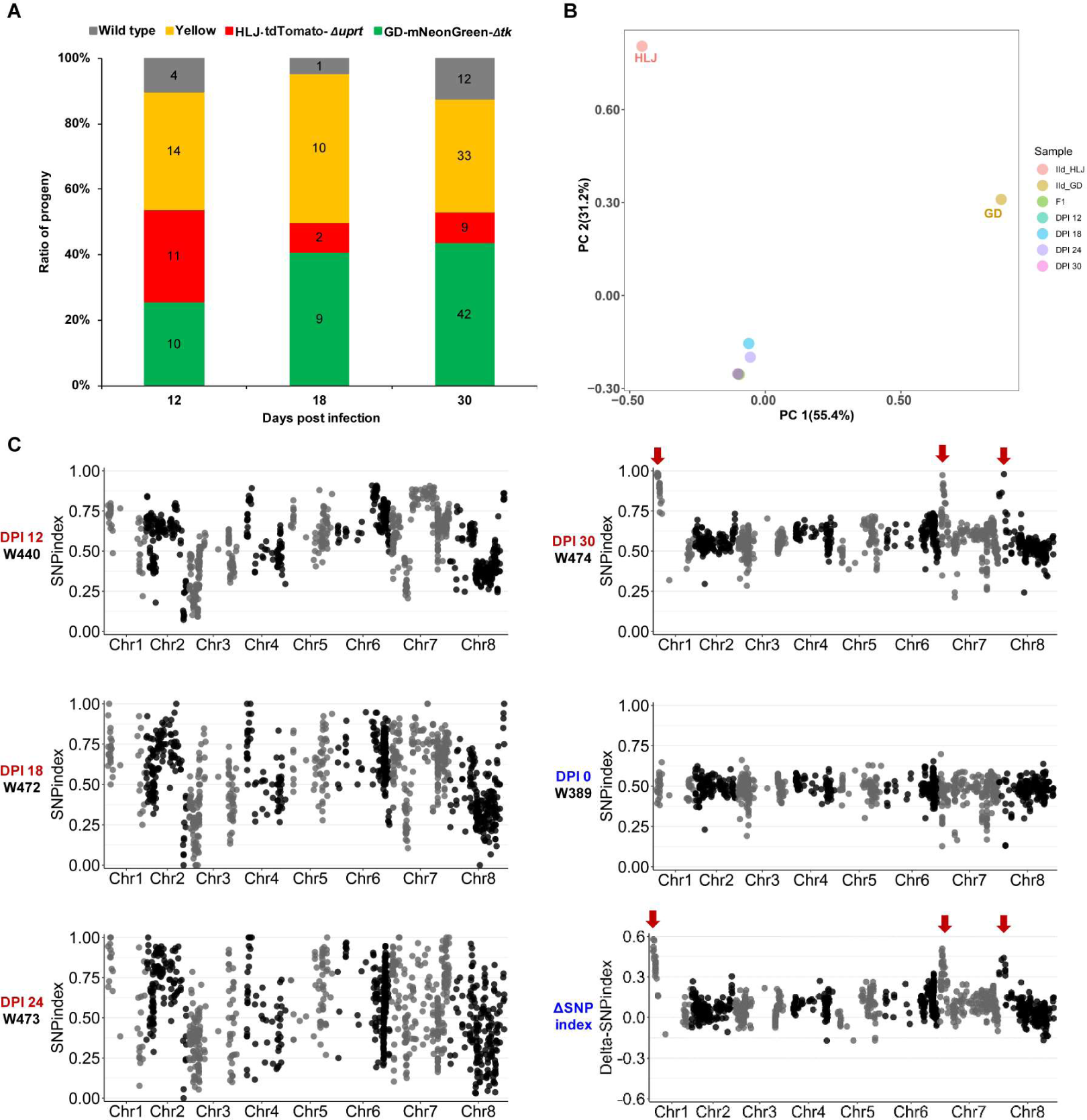
Screening of genes potentially associated with *Cryptosporidium* virulence using BSA in neonatal mice infected with F1 progeny from the second cross of IIdA19G1-GD and IIdA20G1-HLJ treated with paromomycin. (A) The ratio of oocysts of different colors at different time points of infection (n = 7) (B) Principal component analysis (PCA) of WGS data from oocysts collected at different time after infection, where PC1 and PC2 account for variability among samples. (C) Distribution of SNP index of the samples collected throughout the infection.

**Figure S6.**
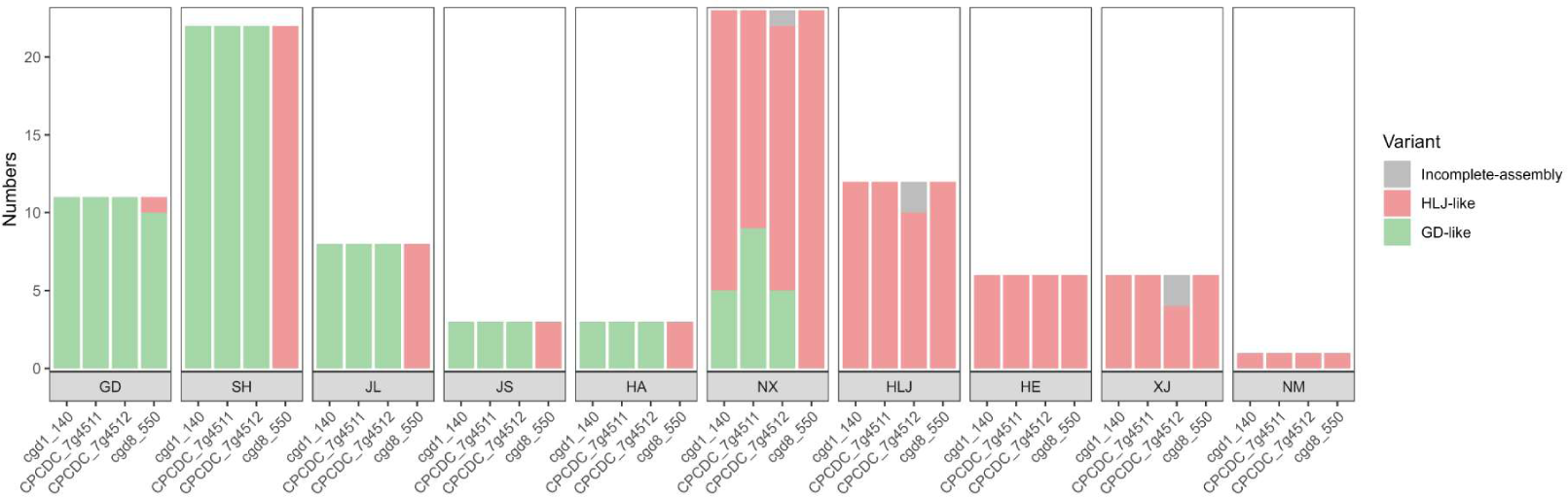
Sequence types at four polymorphic loci potentially associated with virulence among *Cryptosporidium parvum* IId isolates collected from different areas in China. These isolates were whole-genome sequenced, and the sequences of these four genes were extracted from the genome assemblies.^31^

**Figure S7.**
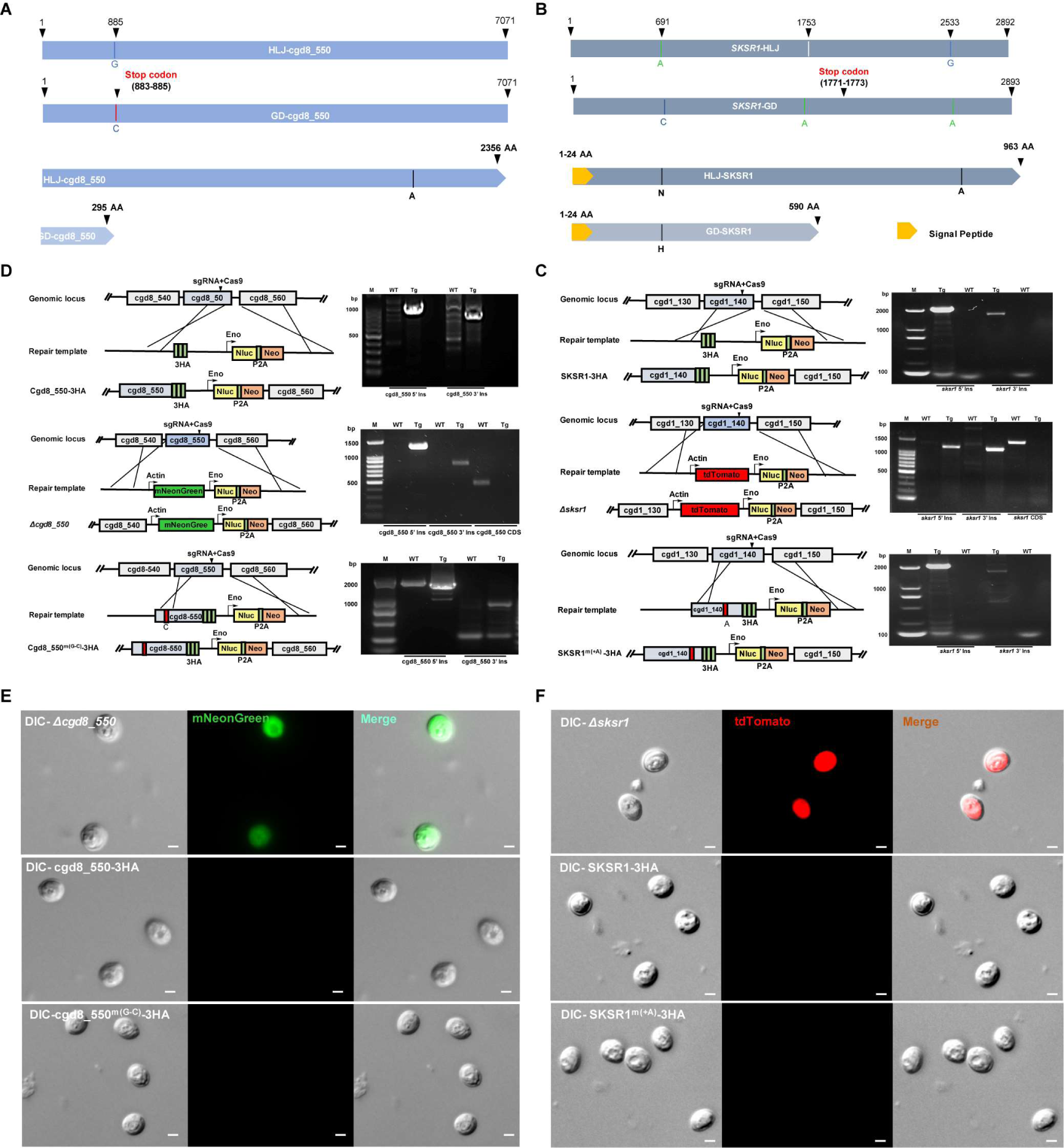
Confirmation of the correct integration of replacement cassettes in different transgenic lines of *Cryptosporidium parvum*. (A and B) Diagrams illustrating the translation changes in the cgd8_550 and *SKSR1* genes caused by base insertions and mutations, respectively. (C and D) PCR confirmation of the correct integration replacement cassettes in different transgenic lines. A schematic map of the modified locus is shown on the left and the corresponding integration PCR gel is shown on the right. The agarose gels show correct 5’ and 3’ integration of the replacement cassette and the absence of a WT band at the modified locus. (E and F) Images of purified oocysts from the *Δcgd8_550* (green) and *Δsksr1* (red).

**Figure S8.**
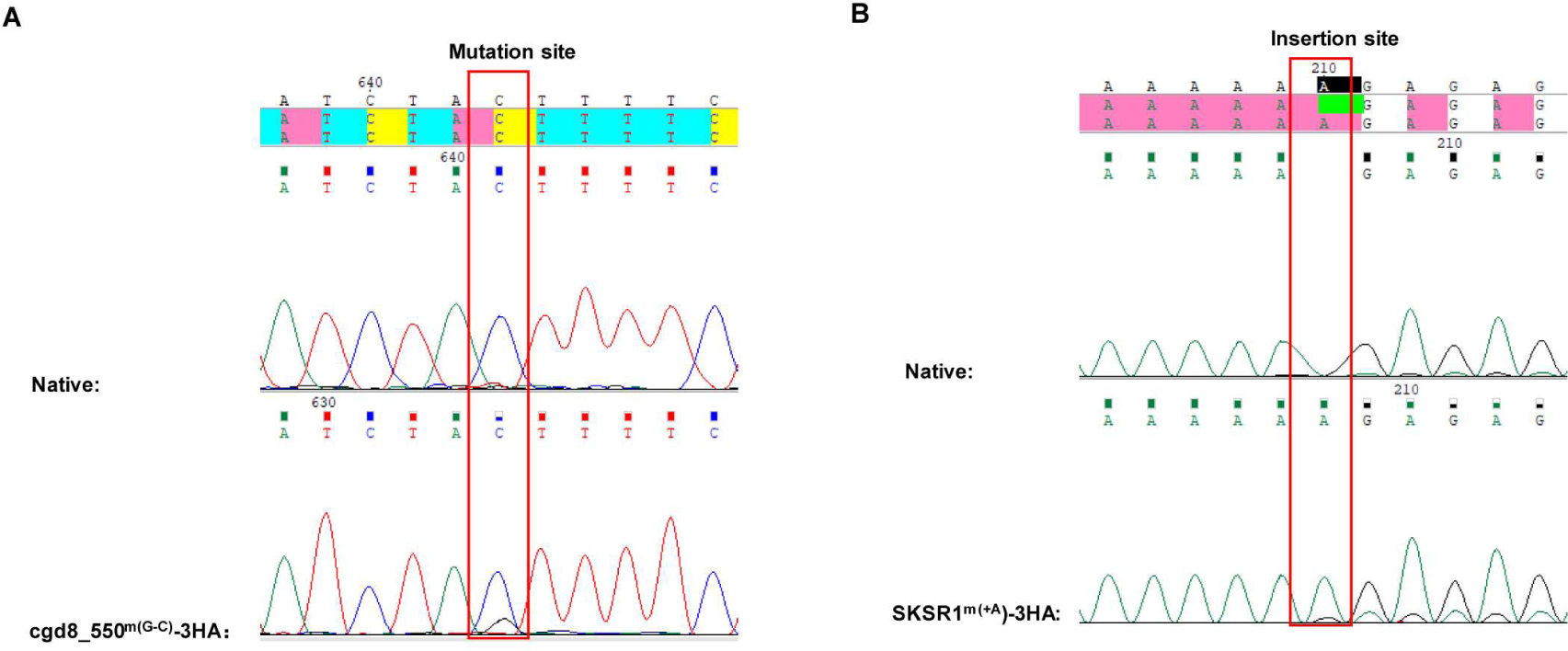
Sequencing electropherograms of PCR products from IIdA20G1-HLJ lines with the mutant and native cgd8_550 and SKSR1 genes. (A) Sequence alignment of PCR products from parasite lines with modified (cgd8_550^m(G–C)^-3HA) and native cgd8_550 gene. (B) Sequence alignment of PCR products from parasite lines with modified (SKSR1^m(+A)^-3HA) and native SKSR1 gene.

**Figure S9.**
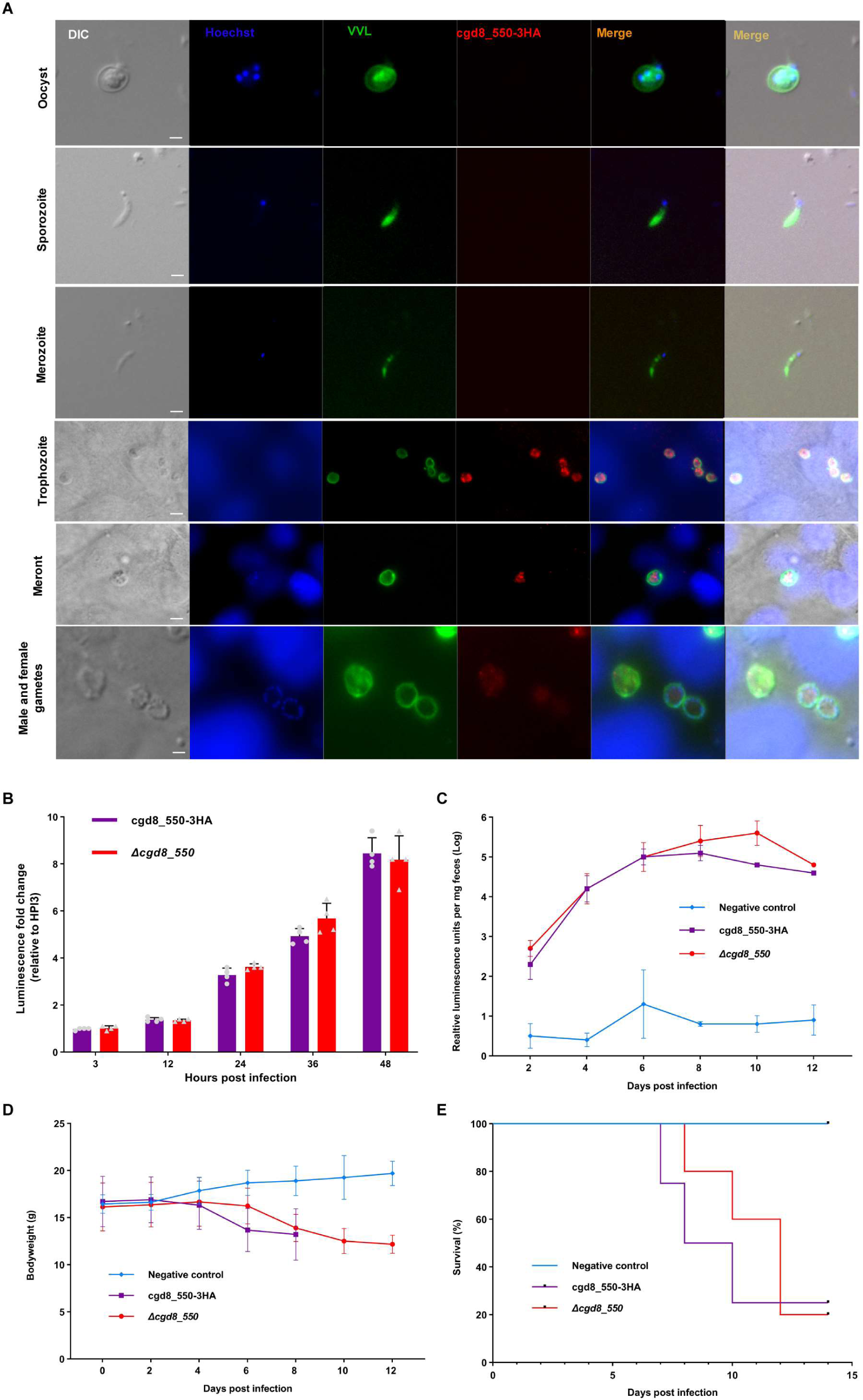
Deletion of the cgd8_550 gene does not affect *Cryptosporidium* growth and pathogenicity. (A) Immunolocalization of cgd8_550 expression. The gene is most highly expressed in trophozoites and meronts. Scale bar is 2 µm. (B) Infection pattern of *Δcgd8_550* and cgd8_550-3HA lines of IIdA20G1-HLJ *in vitro*. There is no difference between the growth of *Δcgd8_550* and cgd8_550-3HA lines. (C) Infection pattern of *Δcgd8_550* and cgd8_550-3HA lines in GKO mice infected with 10,000 oocysts each (n = 5 for each infection group and n = 3 for the uninfected control group). One underweight mouse in the cgd8_550-3HA group died 4 days after infection. (D and E) Body weights and survival curves of mice in the infection study.

**Figure S10.**
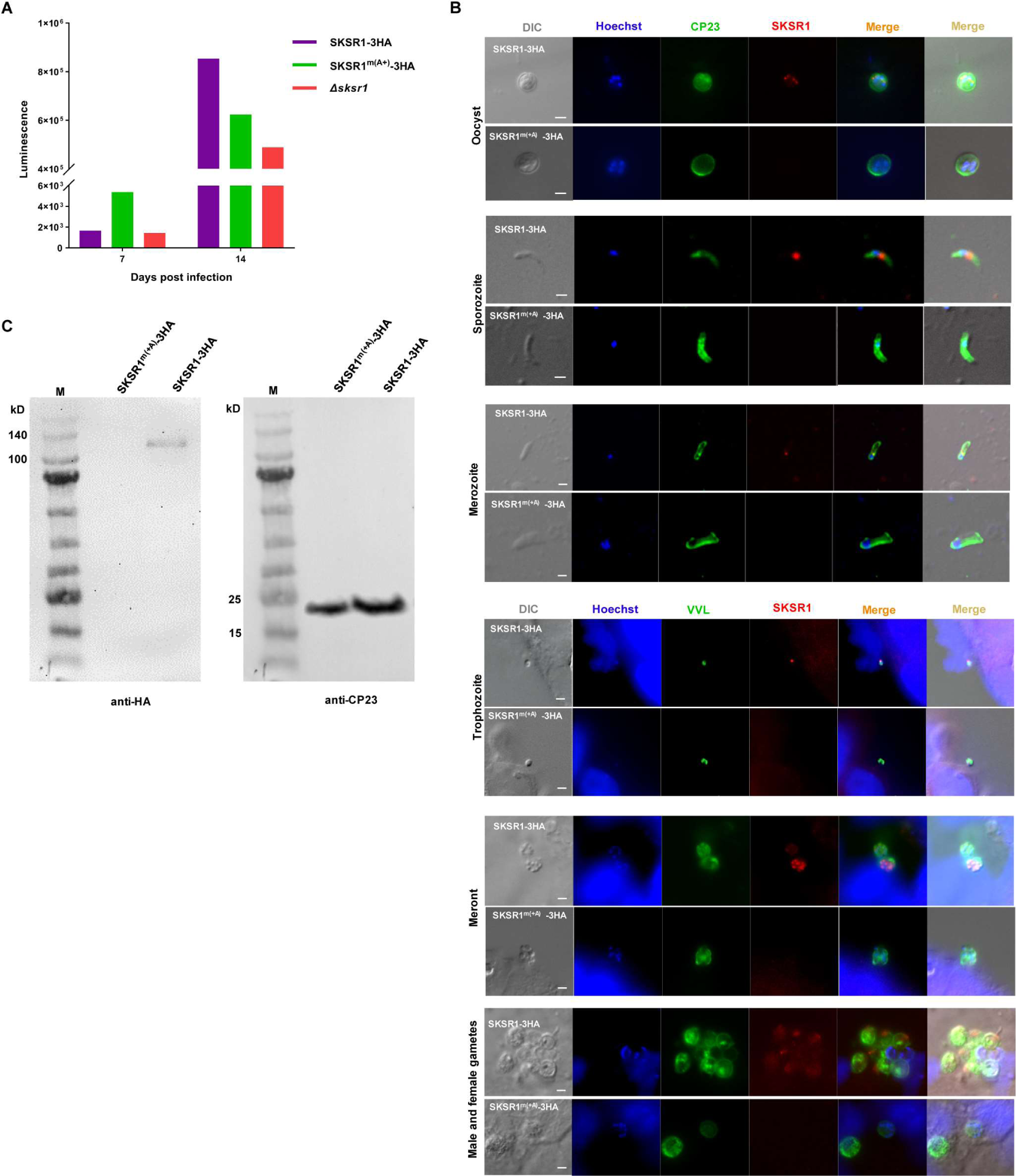
Characteristics of SKSR1 expression in *Cryptosporidium parvum*. (A) Fecal luminescence levels in mice infected with sporozoites transfected with Cas9 guide and repair templates of SKSR1-3HA, *Δsksr1*, and SKSR1^m(+A)-^3HA at DPI 7 and DPI 14. The level was significantly lower in mice infected with *Δsksr1* and SKSR1^m(+A)-^3HA at the peak of infection (DPI 14). (B) Identification of SKSR1-3HA and SKSR1^m(+A)^-3HA expression by IFA. SKSR1 is localized near the nucleus as aggregates (one or two dots) and may be a small dense granule protein, whereas SKSR1^m(+A)^-3HA is not expressed throughout the life cycle. Scale bar 2 µm. (C) Identification of SKSR1-3HA and SKSR1^m(+A)^-3HA expression by Western blot, with the analysis of CP23 expression using a polyclonal antibody as a positive control. The size of SKSR1 in the SKSR1-3HA line was approximately 120 kDa, which is consistent with the expected size. This result confirms that SKSR1 is not expressed in SKSR1^m(+A)^-3HA.

**Figure S11.**
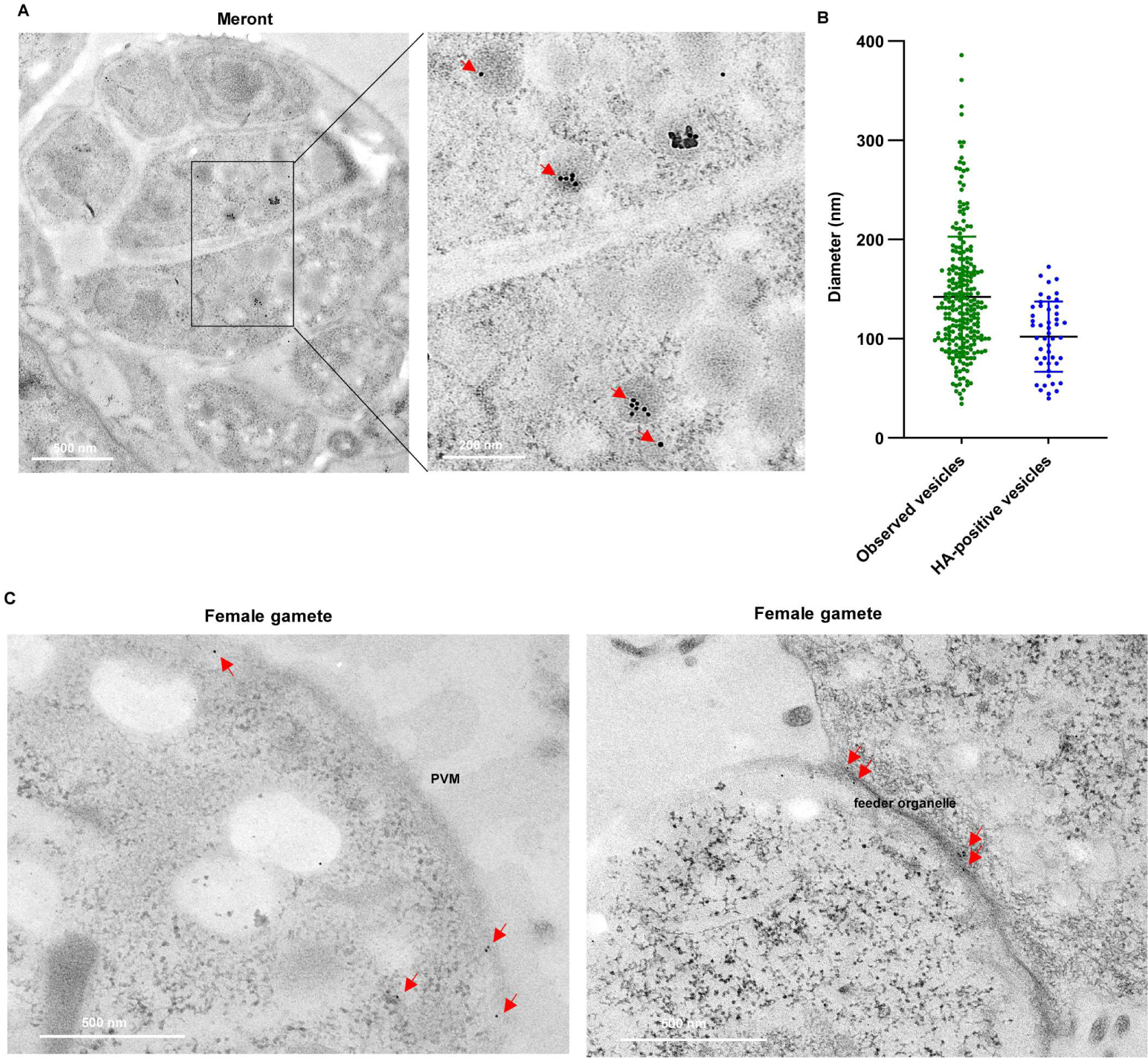
Subcellular localization of SKSR1 by immunoelectron microscopy. (A) Distribution of SKSR1 in small granules of merozoites within a mature meront. (B) Measurement of the size of observed vesicles in IEM images, showing a mean diameter of 102 nm for the HA-positive vesicles. (C) Presence of SKSR1 on the surface (left) and in the feeder organelle (right) of a female gamete.

